# PAD4-mediated histone citrullination contributes to enhanced NETosis in type 2 diabetes but not in type 1 diabetes

**DOI:** 10.64898/2026.06.16.732336

**Authors:** Saijal Gupta, Ankita Priyadarshini, Deepika Sharma, Soumyodeep Das, Samprita Das, Kanika Kalra, Poonam Kumari, Anupam Mittal, Sanjay Bhadada, Parijat Senapati, Sadhan Das

## Abstract

Neutrophil extracellular traps (NETs) are web-like structures released by activated neutrophils through a process known as NETosis, which help immobilize, trap, and eliminate invading pathogens. While NETs play a critical role in host defense, excessive or dysregulated NET formation can contribute to chronic inflammatory diseases such as diabetes and its complications. Elevated levels of NETs and increased expression of its mediator, protein-arginine deiminase type 4 (PAD4), have been reported in diabetes. However, the underlying molecular mechanisms governing the enhanced NETosis in type 1 (T1D) and type 2 diabetes (T2D) remain unclear. Using mouse models and human patient samples, we show that neutrophils undergo enhanced NETosis in T1D and T2D through distinct pathways. We found enhanced NETosis in T1D occurs in a PAD4-independent manner, driven by robust cytosolic ROS production by NADPH oxidase (NOX). This leads to myeloperoxidase activation and chromatin decondensation. We further confirm the PAD4 independent mechanism in neutrophils from STZ-induced PAD4^-/-^ mice. In contrast, in T2D, neutrophils undergo NOX-independent NETosis, which relies on calcium-mediated mitochondrial ROS production, PAD4 activation, and hyper-histone citrullination. Taken together, our findings reveal previously uncharacterized distinct mechanisms of enhanced NETosis in T1D and T2D.

## Introduction

Diabetes mellitus is a highly prevalent disease characterized by hyperglycemia that can result from inadequate insulin secretion by pancreatic islets (T1D) or insulin resistance in major tissues (T2D)^1,2^. Inflammation is a major hallmark of both T1D and T2D, driven by activation of the innate immune system. Neutrophils, key players in the innate immune system, provide the first line of defense against invading pathogens. They neutralize pathogens via various mechanisms, including phagocytosis, degranulation, and a more recently characterized mechanism of releasing neutrophil extracellular traps (NETs) formed through a unique cell death process, termed “NETosis”^3^. NETs are extracellular webs of decondensed chromatin, with granular proteases, that together help neutralize the entrapped pathogen^4^. Although NETs were originally perceived as a protective mechanism, they have now been considered detrimental in various sterile inflammatory conditions, including diabetes and diabetes-associated complications such as atherosclerosis, thrombosis, and impaired wound healing^5–7^. Additionally, the diabetic microenvironment predisposes neutrophils towards enhanced NETosis^8–10^. However, the underlying molecular mechanisms of enhanced NETosis in T1D and T2D remain poorly understood.

Diabetes activates neutrophils to produce more superoxide and cytokines^11,12^. This dysregulated neutrophil response contributes to chronic inflammation and tissue damage, which are commonly observed in diabetic conditions. NETosis can occur via an NADPH oxidase (NOX)-dependent or NOX-independent pathway^13^. NADPH oxidase (NOX)-mediated generation of reactive oxygen species (ROS) activates azurophilic proteins, such as myeloperoxidase (MPO) and neutrophil elastase (NE), leading to chromatin decondensation and the release of NETs^14^. This NOX-dependent mechanism is induced by stimulants such as Phorbol 12-myristate 13-acetate (PMA), bacterial lipopolysaccharide (LPS), or bacterial pathogens^4,13^. Nox-independent NETotic pathway is driven by high intracellular calcium (Ca^2+^) levels rather than NOX2. A key calcium-dependent enzyme, peptidylarginine deiminase 4 (PAD4) gets activated and catalyzes histone citrullination (H3Cit) of the chromatin, leading to its decondensation and expulsion of NETs^15–17^. This NOX-independent mechanism is induced by soluble immune complexes, diverse micro-organisms, and calcium ionophore (CI)^13^. While the role of PAD4 in NETosis is well-studied, its role in enhanced NETosis induced by T1D and T2D remains unclear. Furthermore, whether NOX-mediated signaling cascades are involved in enhanced NETosis in T1D and T2D remains unknown. Understanding the detailed molecular mechanisms underlying T1D- and T2D-mediated NETosis, as well as the mediators involved, can help mitigate the effects of NETosis in diabetes and its associated complications.

In this study, we aimed to investigate the underlying molecular mechanisms of enhanced NETosis in T1D, an autoimmune disorder, and T2D, a metabolic disorder. Using mouse neutrophils and polymorphonuclear neutrophils (PMNs) from individuals with T1D and T2D, we show that neutrophils can be activated by two distinct NETotic pathways. In T1D, neutrophils undergo NOX-dependent NETosis, whereas in T2D, neutrophils undergo calcium-mediated NOX-independent NETosis. The enhanced NETosis in T1D is governed by MPO, while PAD4 plays a critical role in T2D. PAD4-driven NETosis is accompanied by high H3Cit levels in T2D but not in T1D. We further validate PAD4 independence in T1D using STZ-induced PAD4^-/-^ mice. Furthermore, we demonstrate that increased cytosolic ROS production is necessary for NOX-dependent NETosis in T1D. In contrast, enhanced NETosis in T2D neutrophils is driven by mitochondrial ROS, which activates PAD4 and is essential for NET formation. In summary, our data reveal that NET activators, such as PMA, enhance NETosis in T1D via MPO, whereas NET activators, such as CI, increase NETosis in T2D via PAD4 and histone citrullination.

## Results

### NETosis is governed by NOX-dependent and NOX-independent pathways in T1D and T2D, respectively

Elevated NETosis has been observed in the neutrophils of T1D and T2D patients^8,18^. Despite T1D being an autoimmune disorder and T2D being a metabolic disorder, both exhibit increased NETosis. However, the pathways underlying NETosis in T1D and T2D remain to be elucidated. To determine the distinct pathways through which T1D and T2D predispose neutrophils to NETosis, we used the bone-marrow-derived neutrophils (BMDNs) of single high-dose streptozotocin (STZ)-induced C57BL/6 mice neutrophils were either treated with phorbol 12-myristate 13-acetate (PMA) to induce NOX-dependent NETosis or calcium ionomycin (CI) to induce NOX-independent NETosis. NET release induced by these agonists was measured with SYTOX Green fluorescence in NETosis assays (Fig. 1A). Our NETosis assays showed that PMA induces NET formation exclusively in the STZ-induced T1D BMDNs whereas CI induces NET formation in the BMDNs of both STZ-induced T1D mice and in control mice (Fig. 1B). In case of neutrophils from db/+ and db/db mice, we observed that while only CI induces NETs in non-diabetic (db/+) controls, both PMA and CI induce higher levels of NETosis in the neutrophils of diabetic (db/db) mice (Fig.1C). This data indicates a pre-activated state of neutrophils in the db/db mice. To assess the extent of NETosis in human individuals with diabetes, we performed the NETosis assay on neutrophils isolated from the blood of T1D and T2D individuals and their respective healthy controls. Similar to our mouse data, we found that neutrophils from T1D patients showed increased NETosis on PMA treatment compared with those from healthy controls (Fig. 1D). However, PMA treatment did not enhance NETosis in neutrophils from T2D patients compared with those from healthy controls (Fig. 1E). CI treatment, on the other hand, did not induce any NETosis in T1D neutrophils (Fig. 1D), but induced NETosis to a greater extent in T2D neutrophils as compared to healthy controls (Fig.1E). Data from both murine models and human patients indicate that T1D neutrophils are primed to undergo NOX-dependent NETosis induced by PMA, whereas T2D neutrophils are primed for NOX-independent NETosis through CI induction.

**Fig. 1.**
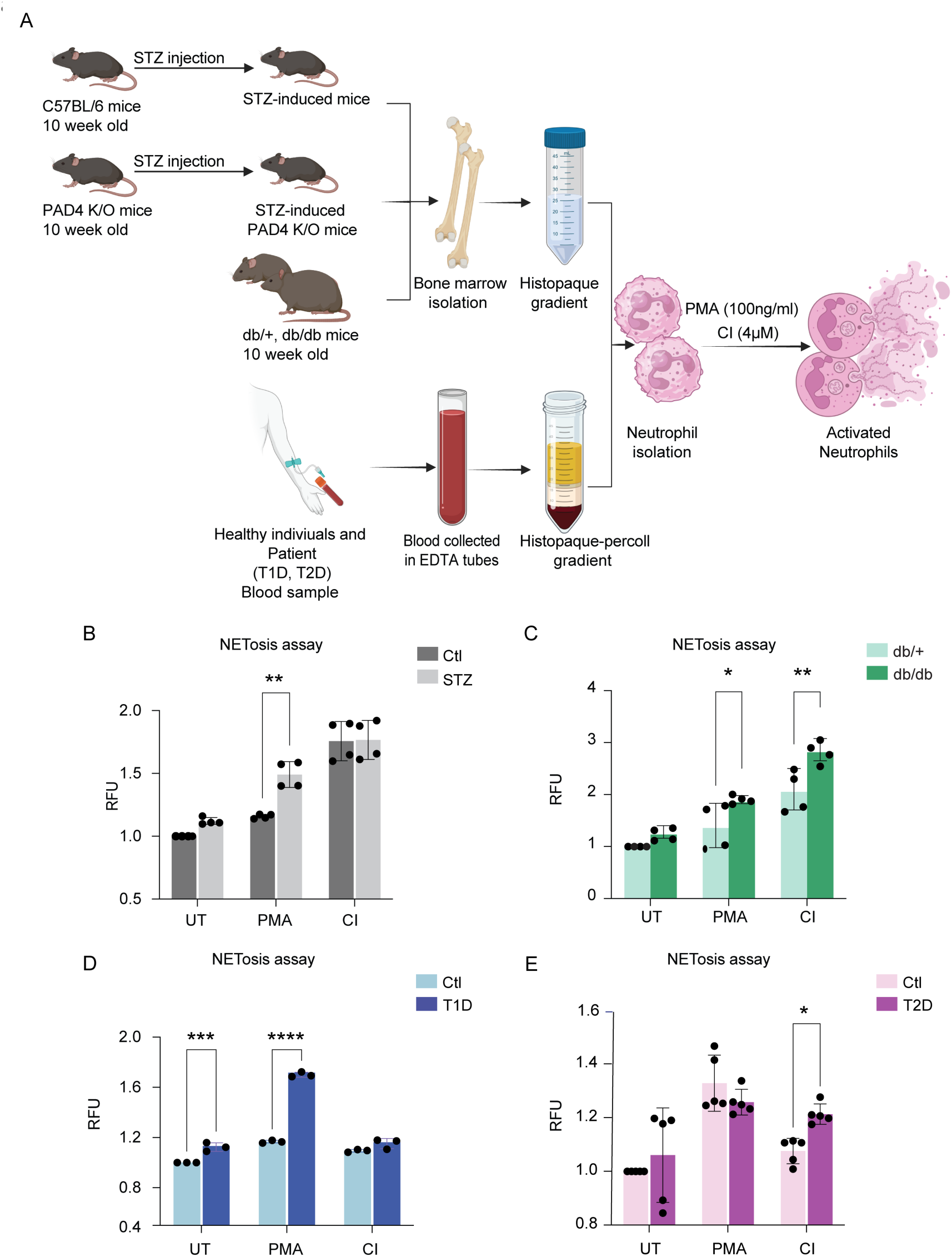
Diabetic microenvironment promotes neutrophil activation and enhances NET formation. (A) Experimental strategy to study NETosis in T1D and T2D mice and humans. (B and C) SYTOX assay showing NET production in PMA-and CI- stimulated neutrophils from STZ-induced mice (T1D) and db/db mice (T2D), respectively (n=4). (D and E) SYTOX assay showing NET production in PMA- and CI-stimulated neutrophils isolated from T1D and T2D human patients’ blood, respectively (n=3-5). \**P* < 0.05, \*\**P* < 0.01, \*\*\**P* < 0.001, \*\*\*\**P*< 0.0001. Data represented as means ± SD.

### Increased PAD4 levels play a key role in promoting NET formation in T2D

PAD4 is a key regulator of NETosis that promotes chromatin decondensation by histone citrullination. Its protein levels are reported to be elevated in neutrophils from diabetic individuals^8,15^. How PAD4 activity is differentially regulated and how it affects enhanced NETosis in T1D and T2D conditions remains an important unanswered question. Since we observed NETosis induction via two distinct pathways in T1D and T2D neutrophils, we tested *Pad4* mRNA expression under these conditions. We found that *Pad4* expression was upregulated under PMA-induced NETosis in neutrophils of STZ-induced mice (Supplementary Fig. 2A) as compared to control mice. However, no change in *Pad4* expression was observed under CI treatment (Supplementary Fig. 2A). Whereas in human T1D patient-derived neutrophils, both PMA- and CI-treatment diabetic controls. In case of neutrophils from db/+ and db/db mice, *Pad4* is upregulated in PMA- and CI-treated db/db neutrophils compared to the control (db/+) (Supplementary Fig. 2C). CI treatment increased the expression of *Pad4* to a greater extent compared to the PMA-treated db/db neutrophils (Supplementary Fig. 2C). Similar to mouse neutrophils, CI augmented the expression of *PAD4* to a greater extent as compared to the PMA-treated T2D neutrophils (Supplementary Fig. 2D).

In addition to mRNA levels, we determined PAD4 protein levels using an immunofluorescence (IF) assay. By analyzing IF data, we found increased PAD4 levels in PMA- and CI-induced neutrophils; moreover, the increase was greater in neutrophils from db/db mice (Fig. 2A and B). Consistent with our data in diabetic mice, we found that PAD4 protein levels are significantly increased in neutrophils from T2D patients treated with PMA and CI compared with those from healthy individuals (Fig. 2C and D). However, there were no significant differences in PAD4 protein levels in the neutrophils of STZ-induced mice (Fig. 2E and F) as compared to those from control mice after PMA- or CI- treatment. Similarly, although PMA and CI increased PAD4 protein levels in neutrophils from T1D patients and healthy individuals, there were no significant differences in PAD4 levels between the control and diabetic groups (Fig. 2G and H). Collectively, these results suggest that enhanced PAD4 levels are likely responsible for enhancing NETosis in T2D but not in T1D.

**Fig. 2.**
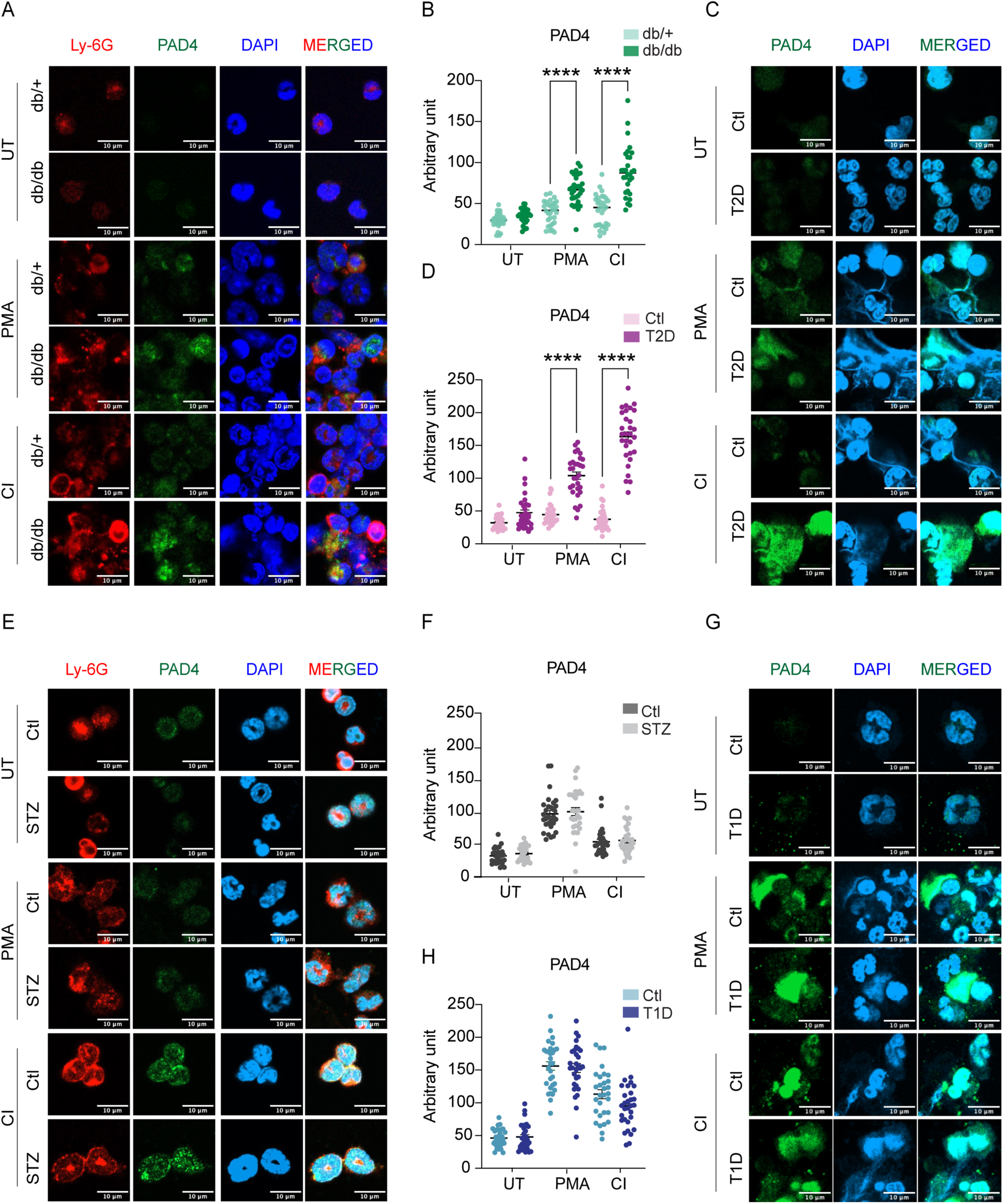
Elevated PAD4 levels play a critical role in promoting NET formation in T2D. (A and E) IF showing PAD4 expression in PMA- and CI-stimulated neutrophils from db/db and STZ-induced mice, respectively, and (B and F) its quantification (n = 30 cells); scale Bar, 10 μm. (C and G) IF showing PAD4 expression in PMA- and CI-stimulated neutrophils from T2D and T1D patients, respectively, and (D and H) its quantification (n = 30 cells); scale bar, 10 μm. \**P* < 0.05, \*\**P* < 0.01, \*\*\**P* < 0.001, \*\*\*\**P*< 0.0001. Data represented as means ± SD.

To further determine the role of PAD4 in diabetes-induced NETosis, we blocked PAD4 function with CI-Amidine, a known pan-PAD inhibitor^19^, and subsequently performed NETosis assays using SYTOX green. We found that CI-Amidine pre-treatment did not result in any significant changes in NETosis in STZ-Induced T1D mice (Fig. 3A). Whereas CI-Amidine pre-treatment blocked NETosis in both PMA and CI-treated neutrophils from db/db as well as db/+ mice (Fig. 3B). Similar to our T1D mice data, we found CI-Amidine treatment had no effect on the PMNs of human T1D patients (Fig. 3C). Whereas CI-Amidine treatment abrogated CI-induced NETosis in the PMNs of T2D patients (Fig. 3D). Together, these results indicate that PAD4 is essential for mediating enhanced NETosis in T2D, as its elevated levels promotes chromatin decondensation and NET formation. In contrast, PAD4 appears to have a lesser role in T1D, suggesting that enhanced NETosis in T1D neutrophils is likely due to the alternate PAD4-independent mechanisms.

**Fig. 3.**
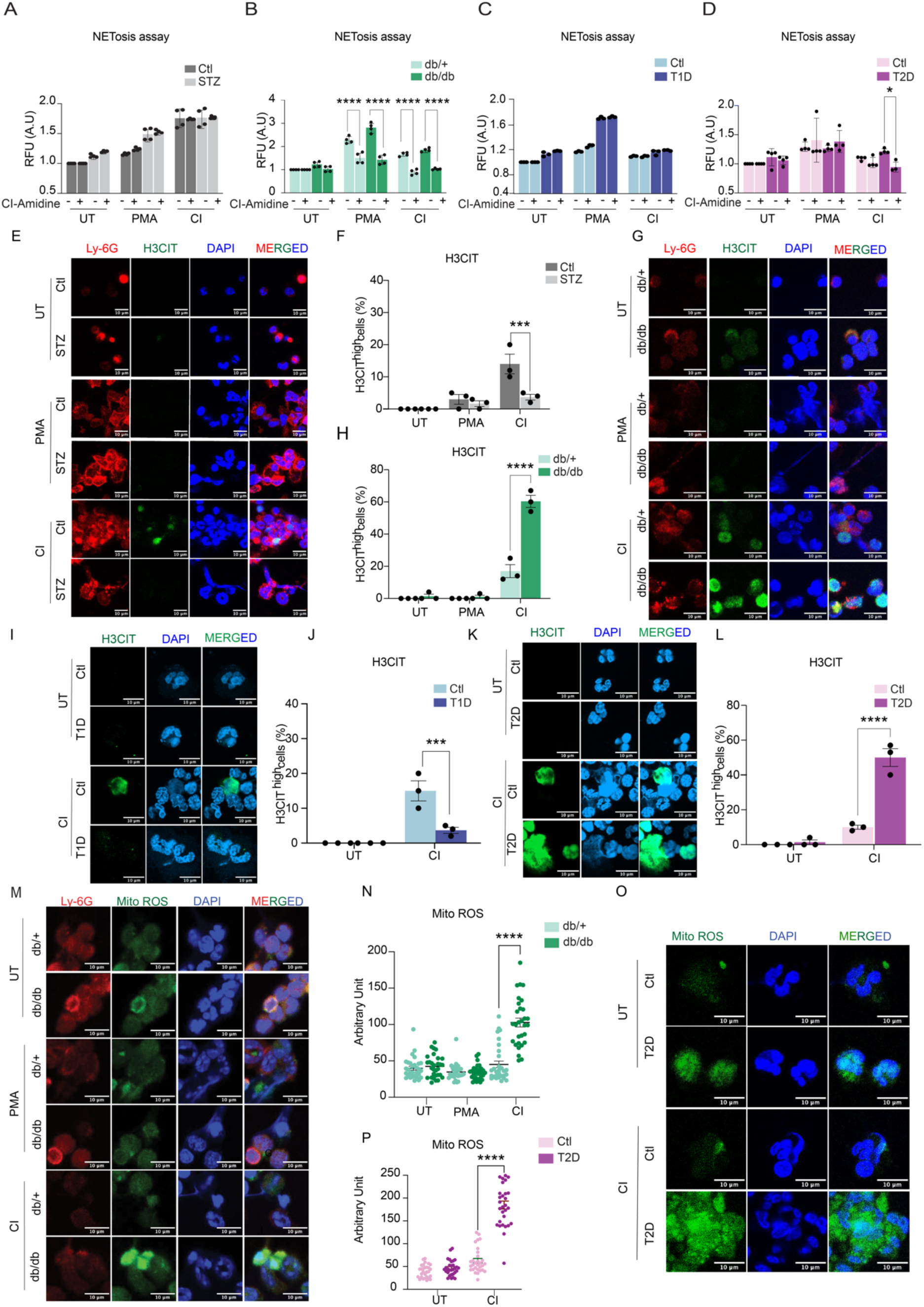
PAD4 activity and histone citrullination are essential for NOX-independent NET formation in T2D. (A and B) SYTOX assay showing effect of Pan-PAD inhibitor (CI-Amidine) in NET formation in T1D and T2D neutrophils from STZ-induced and db/db mice, respectively (n=4). (C and D) SYTOX assay with CI-Amidine, Pan-PAD inhibitor in NET formation in neutrophils from T1D and T2D patient blood samples, respectively (n=3-4). (E and G) IF showing expression of H3CIT in PMA- and CI-stimulated neutrophils from STZ-induced mice and db/db mice, respectively, and (F and H) Percentage of cells that were hypercitrullinated at histone 3 (n=50 cells), scale bar, 10μm. (I and K) IF showing H3CIT level in CI-stimulated neutrophils from T1D and T2D patient blood sample respectively and (J and L) Percentage of cells that were hypercitrullinated at histone 3 (n=50 cells), scale bar, 10μm. (M) IF showing mitochondrial ROS (mtROS) level in PMA- and CI-treated neutrophils from db/db mice and (N) its quantification (n=30 cells). (O) Fluorescence images showing mitochondrial ROS expression in T2D patient sample and (P) its quantification (n =30 cells), scale bar, 10μm. \**P* < 0.05, \*\**P* < 0.01, \*\*\**P* < 0.001, \*\*\*\**P*< 0.0001. Data represented as means ± SD.

### Enhanced NETosis in T2D is mediated by histone citrullination

Since chromatin decondensation is a prerequisite for NET formation^3,15,20^, we sought to assess the roles of mediators other than PAD4 in this process in T1D and T2D. First, we evaluated histone citrullination levels (H3Cit), a PAD4-mediated post-translational modification, using western blot and IF analyses in neutrophils from T1D and T2D mice and in PMNs from T1D and T2D patients. Our western blot data clearly revealed the absence of H3Cit in PMA- and CI-induced neutrophils from T1D mice (Supplementary Fig. 3B), highlighting the PAD4-independent pathway as previously shown. Interestingly, we found increased H3Cit in the CI-treated neutrophils of both db/+ and db/db mice (Supplementary Fig. 3B and C). In human neutrophils, H3Cit levels increased in PMA- and CI-treated PMNs from both control and T1D patients, indicating that diabetic conditions do not have an additional effect on H3Cit (Supplementary Fig. 3D and F). PAD4 levels were elevated even in untreated PMNs from T1D patients, consistent with earlier reports^8^. PMA-treated neutrophils showed an increase in PAD4 levels compared to the untreated PMNs, but there was no further increase under diabetic conditions (Supplementary Fig. 3D and G). In T2D patients, levels of both H3Cit and PAD4 increased in CI-induced PMNs compared with the UT healthy controls (Supplementary Fig. 3E, H, and I). In addition to our western blot data, our IF data also showed that only a few CI-stimulated neutrophils stained positive for H3CIT in control and STZ-induced T1D mice (Fig. 3E and F). However, db/db mice exhibit more H3Cit-positive cells than db/+ mice when NETosis is induced by CI treatment (Fig. 3G and H). Moreover, to ensure human relevance, we also determined H3Cit levels in PMNs from T1D and T2D patients. We found a few H3CIT-positive neutrophils in the CI-treated PMNs of T1D patients (Fig. 3I and J), in contrast to a significantly higher number in the CI-treated PMNs of T2D patients (Fig. 3K and L). For further clarity, we have shown lower-magnification images covering larger fields to illustrate the differences in the number of H3Cit-positive cells in CI-treated control, T1D, and T2D PMNs. A higher number of H3Cit-positive cells is observed only in the T2D group compared to the control and T1D (Supplementary Fig.4A). We also examined the *in vivo* relevance of the histone citrullination in the pancreatic sections of T1D and T2D mice. Our IF results showed higher H3Cit levels in pancreatic sections of diabetic (db/db) mice, but no change in T1D pancreatic sections compared to the respective controls (Supplementary Fig. 4B and C). Together, these data clearly indicate that enhanced histone citrullination during NETosis occurs in T2D, but not in T1D.

**Fig. 4.**
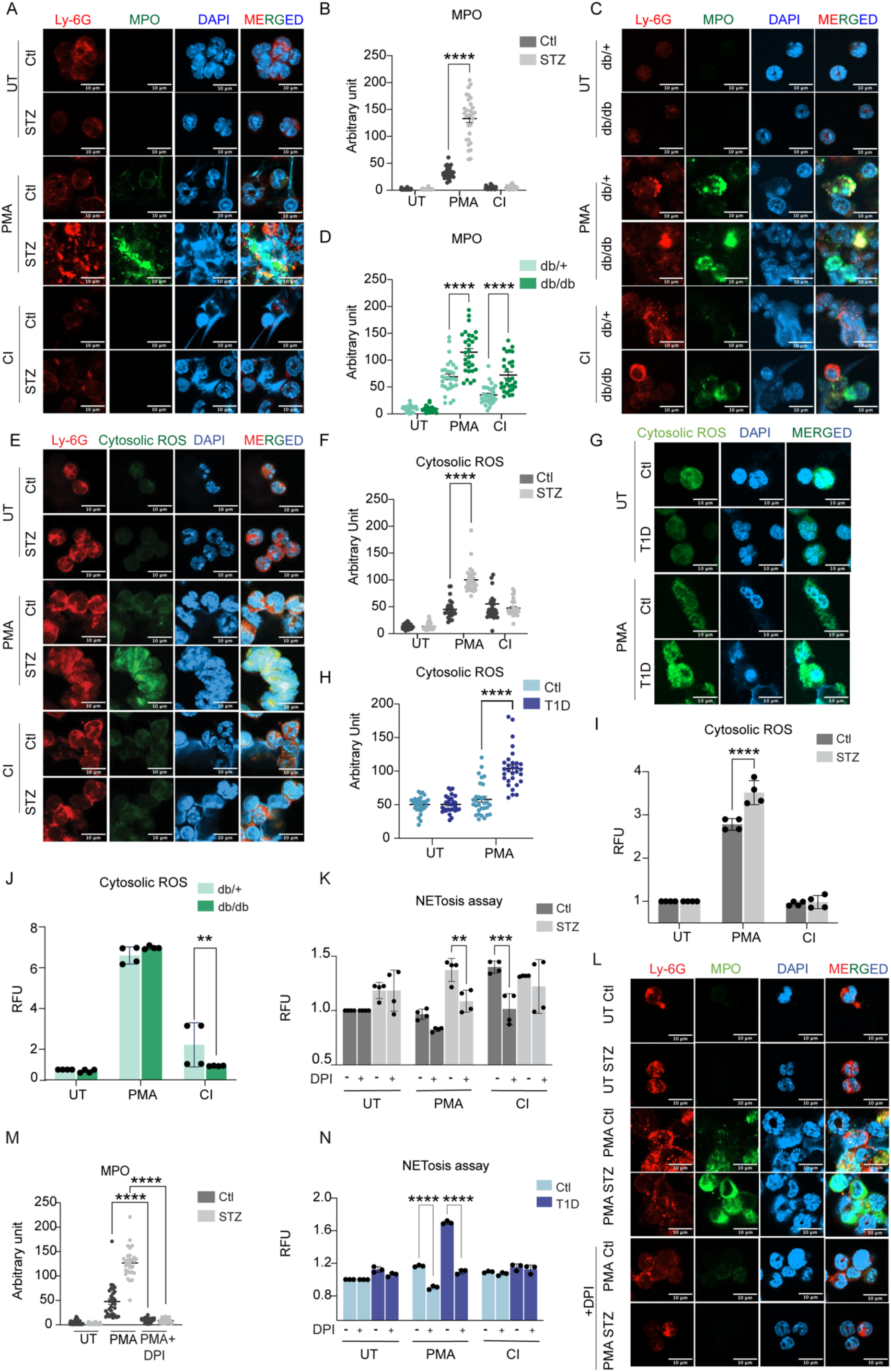
Cytosolic ROS–driven upregulation of MPO enzymatic activity is a key determinant of chromatin decondensation and NETosis in T1D. (A and C) IF showing MPO protein level in PMA- and CI-stimulated neutrophils isolated from STZ-induced and db/db mice, respectively, and (B and D) its quantification (n=30 cells), scale bar, 10μm. (E) IF showing cytosolic ROS level in PMA- and CI-treated neutrophils from STZ-induced mice and (F) its quantification (n=30 cells), scale bar, 10μM. (G) Fluorescence images showing cytosolic ROS level in PMA-treated neutrophils from T1D patients and (H) its quantification (n=30 cells), scale bar, 10μm. (I) DHR123 (Dihydrorhodamine 123) expression showing cytosolic ROS levels in PMA- and CI-treated neutrophils from STZ-induced mice, (n=4). (J) DHR123 expression showing total ROS levels in PMA- and CI-treated neutrophils from db/db mice, (n=4). (K) SYTOX assay showing role of ROS in PMA- and CI-treated neutrophils with ROS inhibitor DPI in NET formation in STZ-induced mice, (n=4). (L) IF showing reduced MPO protein expression upon ROS inhibitor (DPI) in PMA-treated neutrophils from STZ-induced mice and (M) its quantification (n=30 cells), scale bar, 10μm. (N) SYTOX assay showing role of ROS in PMA- and CI-treated neutrophils with ROS inhibitor DPI in NET formation in neutrophils of T1D patients, (n=3). \**P* < 0.05, \*\**P* < 0.01, \*\*\**P* < 0.001, \*\*\*\**P*< 0.0001. Data represented as means ± SD.

### Mitochondrial ROS mediates NOX-independent NET formation in T2D

Our data suggest that increased NETosis in T2D is mediated by PAD4 through a NOX-independent pathway. It is well established that PAD4 requires calcium for its enzymatic activity^16^. Ca²⁺ influx plays a critical role in neutrophil function by modulating mitochondrial ROS production^21^. Therefore, we further explored the role of mitochondrial ROS in calcium-induced NOX-independent NETosis. To determine the production of mitochondrial ROS, we used MitoSOX, a mitochondrial ROS-specific fluorescent dye. Our IF data showed that CI-stimulated neutrophils produced mitochondrial ROS to a greater extent in db/db mice than in control db/+ neutrophils (Fig. 3M and N), an effect not observed with PMA activation. For further clarity, we have also shown images of different fields with results similar to those in Fig. 3M (Supplementary Fig. 5A). However, neutrophils from STZ-induced T1D mice showed reduced mitochondrial ROS production upon PMA stimulation (Supplementary Fig. 5B and C). Moreover, we quantitatively measured mitochondrial ROS using a plate reader assay and found results similar to those of the IF assay (Supplementary Fig. 6A and B). Further, we determined mitochondrial ROS levels in CI-stimulated PMNs from T1D and T2D patients. Consistently, our data demonstrate decreased mitochondrial ROS production in CI-induced PMNs from T1D patients (Supplementary Fig. 6C and D) and increased mitochondrial ROS production in CI-induced PMNs from T2D patients (Fig. 3O and P). Moreover, we observed higher mitochondrial ROS levels in unstimulated diabetic neutrophils than in non-diabetic neutrophils, suggesting that diabetes or an inflammatory microenvironment in T2D enhances NETosis. Thus, T2D may prime neutrophils through increased calcium influx, which elevates mitochondrial ROS, activates PAD4, and ultimately promotes NET formation.

Additionally, to determine whether mitochondrial ROS regulates NOX-independent NETosis in T2D, we used MitoTEMPO, a mitochondrial-targeted antioxidant that scavenges mitochondrial ROS. We treated diabetic neutrophils with MitoTEMPO and performed the SYTOX Green NETosis assay. Interestingly, inhibition of mitochondrial ROS production by MitoTEMPO didn’t result in significant suppression of CI-induced NETosis in diabetic neutrophils isolated from STZ-induced T1D and db/db mice (Supplementary Fig. 6E and F). We examined PAD4 levels after MitoTEMPO treatment and found that inhibiting mitochondrial ROS abrogated PAD4 levels (Supplementary Fig. 6G) but did not prevent NETosis in db/db mice. Therefore, we next asked whether mitochondrial ROS is required for NETosis in diabetic patients. The plate reader assay showed that MitoTEMPO treatment did not alter NET formation in neutrophils from T2D patients (Supplementary Fig. 6H). Overall, our results suggest that mitochondrial ROS production plays a significant role in NOX-independent NETosis in T2D, but it is not essential for this process, suggesting the involvement of additional pathways.

### Enhanced NETosis in T1D is mediated by myeloperoxidase

In addition to PAD4, studies have demonstrated a critical role for granular proteases, including MPO, in chromatin decondensation during NETosis. Upon activation, MPO translocates to the nucleus, where MPO-derived hypochlorous acid (HOCl) promotes histone and DNA degradation, thereby facilitating chromatin decondensation^14,22^. To investigate the underlying molecular mechanisms of chromatin decondensation and NETosis in T1D, we evaluated levels of MPO, a primary granule-resident protein implicated in chromatin opening and NET formation^21,23^.

Our IF data showed significantly higher levels of MPO only in PMA-treated neutrophils from STZ-induced T1D mice (Fig. 4A and B) and in PMNs from T1D patients (Supplementary Fig. 7A and B). No MPO signals were found in the CI-treated neutrophils of T1D mice and humans (Fig. 4A and B; Supplementary Fig. 7A and B). In T2D, we found significantly higher levels of MPO in both CI- and PMA-treated neutrophils from db/db mice compared to those from db/+ mice (Fig. 4C and D) and in PMNs from T2D patients (Supplementary Fig. 7C and D). Moreover, we found higher levels of MPO in PMA-treated neutrophils as compared to the CI-treated neutrophils in both T2D mice and humans. Collectively, these findings indicate that MPO is a critical mediator of chromatin decondensation and NETosis in T1D, whereas in T2D, both histone citrullination and MPO are key drivers of chromatin relaxation and NET formation.

### Increased cytosolic ROS leads to NOX-Dependent NETosis in T1D

Our results support that T1D primes neutrophils for NET formation in a NOX-dependent manner. Since the generation of reactive oxygen species (ROS) by NOX2 is crucial for the induction of NOX-dependent NETosis^24^, we first assessed Nox2 mRNA expression upon NETosis induction. RT-qPCR results showed expression of Nox2 is upregulated in both PMA- and CI-treated neutrophils of STZ-induced T1D mice compared to the control mice (Supplementary Fig. 8A). However, no changes in Nox2 expression were observed upon PMA- and CI-treatment in db/db mice (Supplementary Fig. 8B). Then, we measured the production of cytosolic ROS in the PMA- and CI-treated neutrophils of T1D and T2D mice. We used Dihydrorhodamine123 (DHR123), a cell-permeable fluorogenic dye that is oxidized by ROS to form a fluorescent product. Our IF data revealed significantly increased cytosolic ROS production only in PMA-induced neutrophils from T1D mice compared with controls (Fig. 4E and F). In addition, to assess human relevance, we conducted fluorescence imaging using PMA-treated PMNs from T1D patients and observed a significant increase in cytosolic ROS production (Fig. 4G and H). We also found that cytosolic ROS production did not change in PMA-induced neutrophils of T2D mice and T2D patients. Furthermore, there was a decrease in cytosolic ROS production in CI-induced diabetic neutrophils (Supplementary Fig. 9A to D). Further, we confirmed our results by performing a plate reader assay with DHR123, where we found PMA-treated neutrophils from STZ-treated T1D mice indeed had increased cytosolic ROS whereas CI-treated neutrophils showed no change in cytosolic ROS levels (Fig. 4I). In neutrophils from T2D mice, a decrease in cytosolic ROS was observed after CI treatment (Fig. 4J), indicating a less prominent role for cytosolic ROS in NOX-independent NETosis in T2D.

To further assess whether NOX2 is necessary for diabetes-primed NETosis, we performed a SYTOX green NETosis assay with or without Diphenyleneiodonium chloride (DPI), a potent inhibitor of NOX (NADPH-oxidase), which is known to reduce the production of NOX-generated ROS. Our results showed that DPI treatment abrogates PMA-induced NET formation in the STZ-induced T1D mice (Fig. 4K). In addition, IF data showed that DPI treatment significantly decreased MPO levels during PMA-stimulated NET formation in STZ-induced T1D mice (Fig. 4L and M). Moreover, we found that DPI decreased PMA-induced NETosis in PMNs from T1D patients (Fig. 4N). Together, our data strongly support the notion that NETosis in T1D follows a NOX-dependent pathway, in which NOX-mediated cytosolic ROS acts as a critical factor in MPO activation, chromatin decondensation, and NET formation. Consistent with this, pharmacological inhibition of NOX2 suppresses ROS production and subsequently inhibits NETosis.

Additionally, we recapitulated our single high-dose T1D (200mg/kg: High-STZ-induced) data in five consecutive low-dose STZ-induced T1D (50mg/kg: Low-STZ-induced) mice^25^. Consistent to the high-STZ-induced mice, we found enhanced NETosis in the PMA-treated low-STZ-induced T1D mice compared to the control (Supplementary Fig. 10A) as shown in Fig. 1B (Supplementary Fig. 10A vs Fig.1B). Similarly, we found increased expression of *Pad4* at mRNA level only in the PMA-treated low-STZ-induced T1D mice (Supplementary Fig. 10B vs Supplementary Fig. 2A). Further we found no change in the PAD4 protein levels in the PMA-and CI-treated low-STZ-induced T1D mice (Supplementary Fig. 10C vs Fig. 2E). In addition, we did not find change in the H3Cit levels in the CI-treated low-STZ-induced T1D mice (Supplementary Fig. 10D vs Fig. 3E). CI-amidine treatment did not have any effect on the production of NETs in low-STZ-induced (Supplementary Fig. 10E vs Fig. 3A). Furthermore, we found higher production of ROS in the PMA-treated neutrophils of low-STZ-induced T1D mice (Supplementary Fig. 10F). DPI treatment led to the reduction of NET production in the low-STZ-induced neutrophils (Supplementary Fig. 10G). Collectively, these data clearly suggest both high- and low-STZ-induced T1D mice exhibit a similar mode of action for enhanced NETosis in T1D and differ from T2D-mediated enhanced NETosis.

### Enhanced NETosis in T1D occurs in a PAD4-independent manner

To confirm that T1D follows NETosis in a PAD4-independent manner, we used PAD4^-/-^ mice. First, we injected five consecutive doses of STZ in PAD4^-/-^ mice for the development of T1D in PAD4^-/-^ mice (Supplementary Fig. 11A). Our data clearly showed that STZ-induced PAD4^-/-^ mice has significant lower body weight compared to the control and increased blood glucose levels compared to the control, respectively (Supplementary Fig. 11B and C). Further, H&E and IF data revealed a reduction in the insulin-producing β cells in STZ-induced PAD4^-/-^ mice (Supplementary Fig. 11D and E). Upon successful development of STZ-induced T1D in PAD4^-/-^ mice, we performed an IF assay with MPO in the BMDNs isolated from PAD4^-/-^ mice and STZ-induced T1D in PAD4^-/-^ mice with or without PMA and CI stimulation. Our IF data showed higher levels of MPO in the PMA-treated STZ-induced T1D PAD4^-/-^ mice (Fig. 5A and B), suggesting an important role of MPO in the formation of NETs in T1D, as is evident in STZ-induced T1D mice (Fig. 4A) and T1D patients (Supplementary Fig. 7A). Further, our SYTOX Green NETosis assay showed increased NETosis only in the PMA-treated STZ-induced T1D PAD4^-/-^ mice (Fig. 5C), suggesting NETosis follows NOX-dependent pathways in T1D. We also showed that DPI treatment attenuates NET formation in PMA-treated STZ-induced T1D PAD4^-/-^ mice (Fig. 5D).

**Fig. 5.**
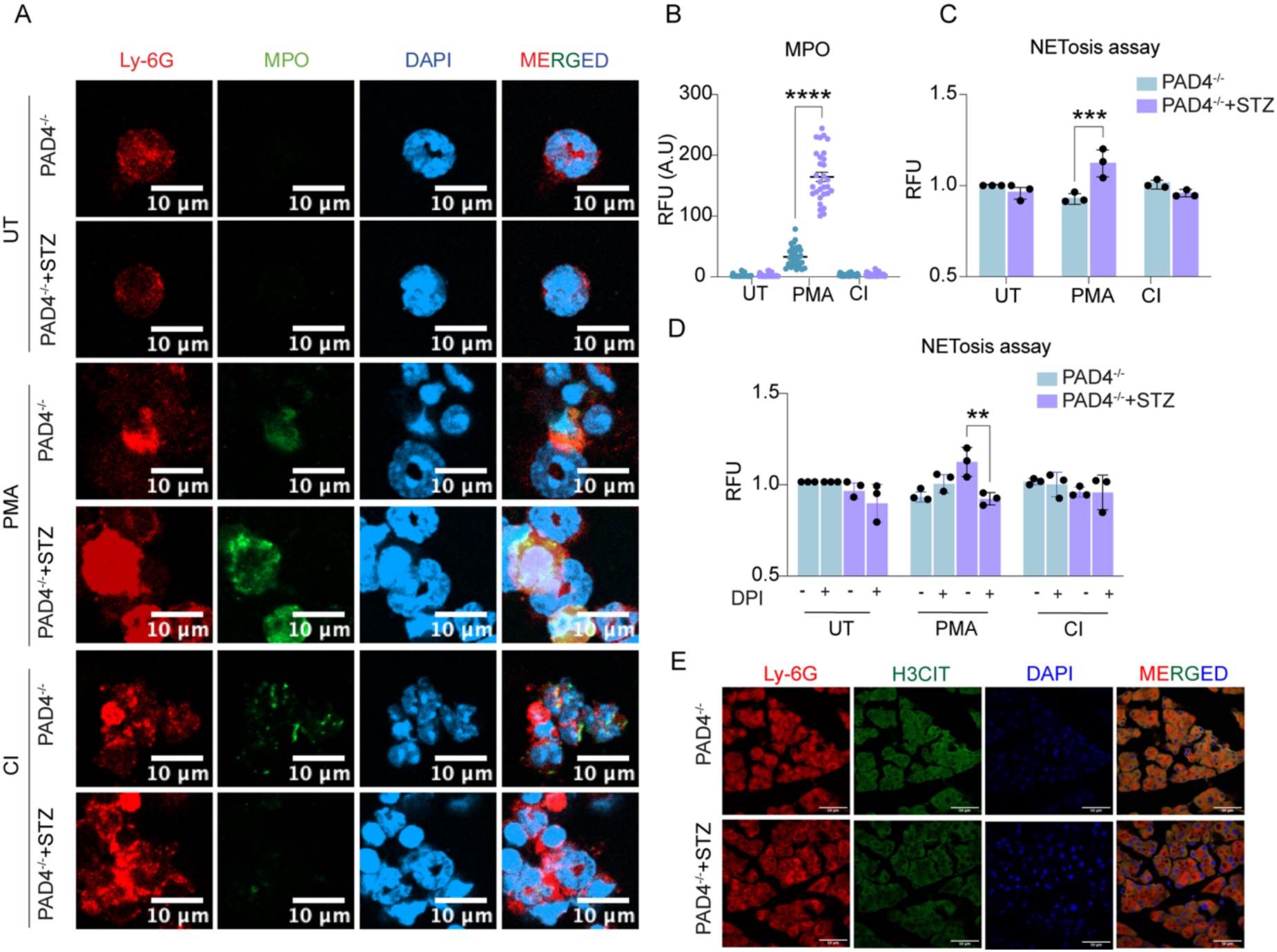
T1D reprograms neutrophils toward PAD4-independent, NOX-dependent NETosis, as demonstrated in STZ-induced PAD4-deficient mice. (A) IF showing MPO expression in neutrophils isolated from STZ-induced PAD4^-/-^ mice and (B) its quantification (n =30 cells), scale bar, 10μm. (C) SYTOX assay showing NET production in PMA- and CI-stimulated neutrophils isolated from STZ-induced PAD4^-/-^mice (n=3). (D) SYTOX assay showing role of ROS in PMA- and CI-treated neutrophils with ROS inhibitor DPI in NET formation in neutrophils of STZ-induced PAD4^-/-^ mice (n=3). (E) Immunofluorescence image showing H3CIT abundance in pancreatic sections isolated from STZ-treated PAD4^-/-^ mice. \**P* < 0.05, \*\**P* < 0.01, \*\*\**P* < 0.001, \*\*\*\**P*< 0.0001. Data represented as means ± SD.

Interestingly, our *in vivo* study revealed no changes in the H3Cit levels in the pancreatic section of the STZ-induced PAD4^-/-^ mice (Fig. 5E), indicating no involvement of histone citrullination in T1D NETosis. Altogether, these data suggest that T1D diabetic neutrophils undergo PAD4-independent and NOX-dependent NETosis.

## Discussion

The present study demonstrates discrete mechanisms underlying the elevated NETosis in T1D and T2D. By employing well-known NET activators, PMA and CI, we show that PMA preferentially induces NET formation in a NOX-dependent manner in STZ-induced T1D mice, whereas CI enhances NETosis in a NOX-independent manner in db/db (T2D) mice. Notably, these findings were consistent across human PMNs derived from T1D and T2D patients, suggesting that the divergent NETotic pathways observed in murine models are translationally relevant to humans. Currently available therapeutic approaches aim to mitigate NET-driven pathology by pharmacologically inhibiting NET formation or the enzymatic degradation of NETs. However, due to a lack of understanding of the underlying NET biology in T1D and T2D, there are no NET-targeting drugs tailored for diabetes or its associated complications. Herein, the study delineates two distinct NETotic pathways in T1D and T2D, which can help inform complication-specific interventions and the precise design of NET-targeted therapies.

T1D primarily arises from the destruction of pancreatic β cells, leading to insulin deficiency, whereas T2D is characterized by β-cell dysfunction and insulin resistance. Despite their distinct etiologies, both forms of diabetes exhibit chronic, low-grade inflammation, in which immune cells, particularly neutrophils, are central players^26–29^. Accumulating evidence suggests that neutrophils in diabetic conditions exist in a primed state and are more susceptible to NET formation^8^. Although NETosis is a protective mechanism against invading pathogens, excessive or dysregulated NET release contributes to chronic inflammation and tissue damage, as seen in diseases such as sepsis and diabetes^30–32^. NETosis is a multifaceted process orchestrated by a series of interconnected molecular events mainly driven by PAD4. PAD4 is a calcium-dependent enzyme responsible for histone citrullination and chromatin decondensation, contributing to expulsion of NETs^15,33,34^. Therefore, PAD4-mediated histone citrullination is widely recognized as a marker of NETosis. Nevertheless, extensive research in this area has revealed that diverse stimuli activate distinct NETotic pathways that do not necessarily depend on PAD4 activation^34,35^. In agreement with this notion, we observed a striking divergence in T1D and T2D NETotic pathways regarding PAD4-mediated citrullination and PMA/CI-induced NETosis. We report elevated PAD4 levels and histone hypercitrullination in T2D mice and patients upon CI stimulation. In contrast, there was no change in the PAD4 or citrullination levels in T1D mice and patients following NET induction. Further, pharmacological inhibition of PAD activity by CI-amidine markedly inhibited NET formation in T2D neutrophils with no significant effect in T1D NET formation. Our data strongly indicate that PAD4-mediated histone citrullination contributes to NETosis specifically in T2D and not in T1D.

To further dissect the mechanism of PAD4-mediated NETosis in T2D, we examined the upstream pathways regulating PAD4 activation. Several studies have shown that the intracellular ROS production is increased in a diabetic microenvironment, leading to PAD activation and NET formation^11,36^. It is further reported that CI stimulates the mitochondrial ROS production in a NOX-independent manner, while PMA induces the cytosolic ROS generation in a NOX-dependent manner^13,19^. In this study, we found higher mitochondrial ROS production in CI-treated neutrophils of both T2D mice and patients, further supporting the notion that NET formation in T2D follows a NOX-independent pathway. Surprisingly, we found that mitochondrial ROS depletion reduced PAD4 expression without affecting NETosis (Supplementary Fig. 6 F and G). However, we found significant abrogation of NETosis upon treatment with pan-PAD inhibitors in T2D neutrophils, suggesting a role for other PAD-family enzymes beyond PAD4. For instance, studies have shown that selective inhibition of PAD2 can reduce H3Cit levels in NETotic neutrophils. Interestingly, our gene expression data also showed increased *Pad2* expression in NETosis-induced neutrophils of T2D mice (Supplementary Fig. 12A and B). Collectively, these data suggest a possible role for other PAD family proteins in NOX-independent NETosis in T2D, which can be explored in future studies.

On the contrary, PMA-induced NETosis in T1D is regulated through a NOX-dependent pathway. While the PMA-induced NETosis has been reported by us and others, the present findings provide additional mechanistic insight driving this process. We observed robust cytosolic ROS production during PMA-induced NETosis in both T1D mice and patients, indicating NOX2-mediated ROS is essential for NOX-dependent NETosis. Furthermore, pharmacological inhibition of NOX-derived ROS using DPI significantly reduced PMA-induced NET formation in T1D mice and patients, indicating that cytosolic ROS is indispensable for NETosis. In parallel, studies have also demonstrated that cytosolic ROS induces the release of MPO from the azurophilic granules, driving chromatin decondensation and mediating NET formation^23^. Consistent with these observations, our data also showed significantly higher levels of MPO in PMA-induced neutrophils from T1D patients compared with controls, validating MPO as an important component of NETosis in T1D. Taken together, these findings suggest that enhanced cytosolic ROS production and increased MPO release are key drivers of PMA-induced NETosis in T1D and further highlight the mechanistic differences between NETosis pathways in T1D and T2D.

To further confirm PAD4-independent NETosis in T1D and to consolidate mechanistic details, we generated STZ-induced PAD4^-/-^ mice. STZ-induced T1D PAD4^-/-^ mice showed increased NETosis and MPO levels in the PMA-treated condition as compared to the control. Additionally, inhibiting cytosolic ROS production in STZ-induced T1D PAD4^-/-^ mice attenuated NOX-mediated NET formation, further confirming NETosis in T1D does not depend on PAD4 but on cytosolic ROS. *In vivo* studies also revealed no changes in the H3Cit levels in the pancreatic section of the STZ-induced PAD4^-/-^ mice (Fig. 5E). Taken together, these data suggest that T1D aggravates NET formation via activation of NOX2, increased production of ROS, and higher MPO activity, thereby promoting NOX-dependent NET formation.

In summary, our study demonstrated that T1D primes neutrophils via a PMA-induced, NOX-dependent pathway, leading to NET production, whereas T2D promotes NETosis through a CI-induced, NOX-independent pathway. We further show that the NOX-independent NETosis pathway in T2D requires PAD4 activation and histone citrullination, unlike NETosis in T1D. Together, these results provide important mechanistic insights into the divergent pathways of NETosis in T1D and T2D neutrophils (Fig. 6). Moreover, distinguishing the mechanistic differences in NETosis between T1D and T2D offers the potential to identify disease-specific interventions, enabling more precise and effective therapeutic approaches for diabetes and diabetes associated complications.

**Fig. 6.**
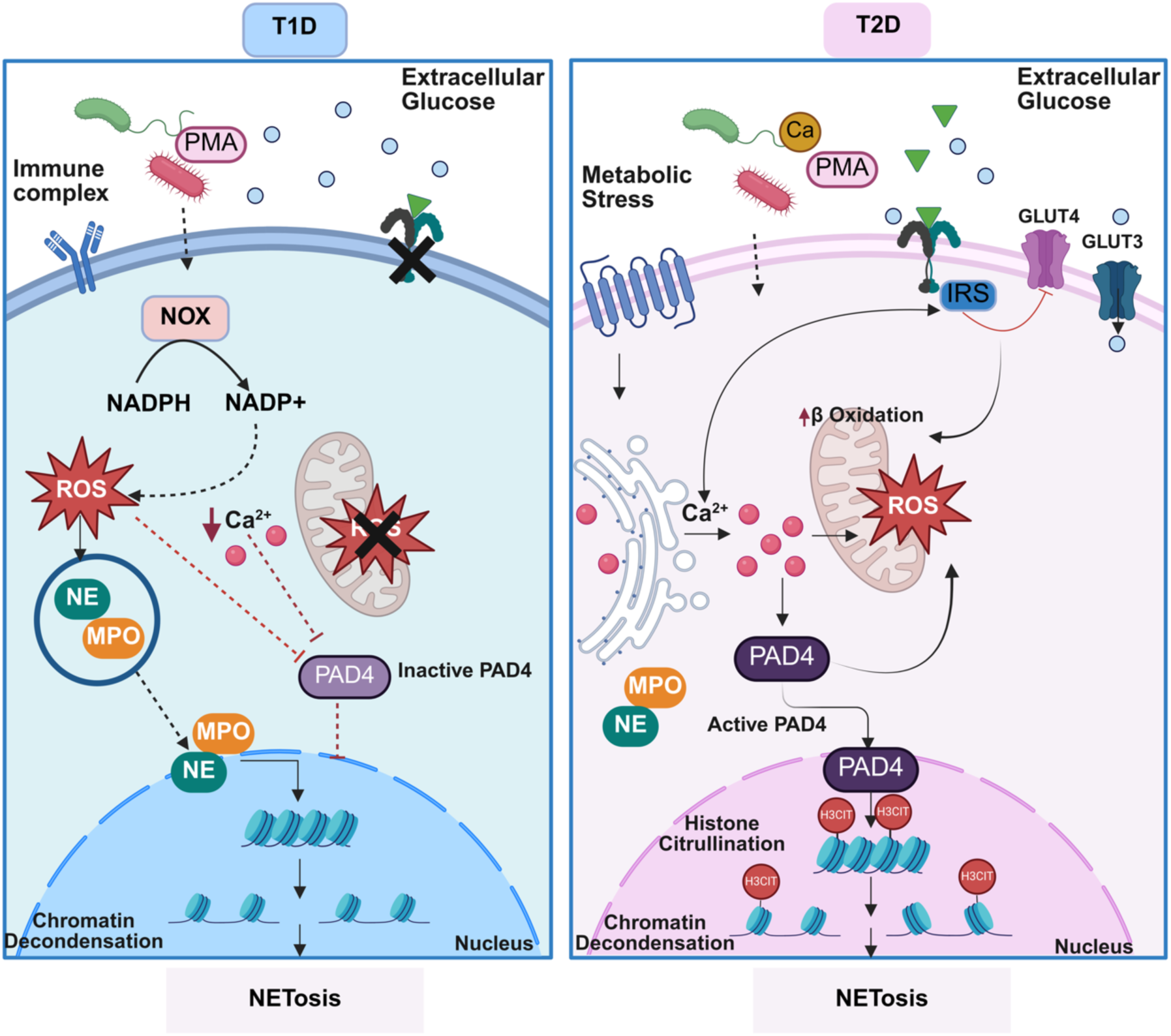
Graphical representation depicting distinct pathways of enhanced NETosis in T1D and T2D. In T1D, Immune complexes, PMA, and LPS activate the NOX gene, an important component of NOX-dependent NETosis, leading to ROS generation in the neutrophils. The cytosolic ROS further activates granular proteases NE and MPO, which migrate to nuclei and lead to chromatin opening. Whereas in T2D, NETosis follows NOX-independent pathway. Metabolic stress and crystals activate neutrophils, leading to calcium efflux and mitochondrial ROS production. The increased calcium concentration activates PAD4, which migrates to the nuclei, leading to histone citrullination and chromatin opening.

## Methods

### T1D Mice and T2D mice

All animal experiments were performed in accordance with the recommendations of the Institutional Animal Ethics Committee (IAEC) at IISER-Mohali. Diabetic (db/db) and C57BL/6 mice were purchased from the Jackson Laboratory (Bar Harbor, ME) and IISER-Mohali, respectively. Mice were housed in a pathogen-free animal facility and were fed a standard chow diet. Male mice were used for all the experiments. For studying T1D, mice were rendered diabetic by either a single high dose or five consecutive low doses of streptozotocin (STZ). C57BL/6 mice received a single high dose of STZ (200 mg/kg) prepared in sodium citrate buffer (Supplementary Fig. 1A). Pancreas were collected, and pancreatic islets were examined by H&E staining and immunohistochemistry with Insulin and DAPI counterstaining (Supplementary Fig. 1D, E). Mice with blood glucose levels exceeding 300 mg/dL were classified as diabetic and used for subsequent experiments (Supplementary Fig. 1B). In addition, T1D was also induced by five consecutive low-doses of STZ injection (Supplementary Fig. 1A). Briefly, 10-week-old male C57BL/6 mice were fasted for 6 hours and then injected intraperitoneally with either vehicle or STZ (50 mg/kg/day) for five consecutive days. Fasting blood glucose levels and body weight were monitored throughout the study (Supplementary Fig. 1F and G). We also examined pancreatic islets by H&E and immunohistochemistry (Supplementary Fig. 1H and I). Mice with fasting blood glucose levels above 300 mg/dL were considered diabetic and used for further experiments.

For studying T2D, db/db mice (BKS.Cg-m+/+leprdb/J) and age-matched non-diabetic littermate db/+ mice were used for the experiments. Body weight and blood glucose levels for both db/+ and db/db mice were measured before conducting the experiments (Supplementary Fig.1J and 1K).

### Isolation of neutrophil and induction of NETosis

Bone marrow cells from the femur and tibia of T1D and T2D mice were isolated. Cells were collected in RPMI 1640 with 10% FBS, 1% Pen/Strep, and RBCs were removed by hypotonic lysis with 0.2% NaCl. Neutrophils were isolated by the Histopaque density gradient method by layering Histopaque-1077 (density: 1.077 g/mL, Sigma-Aldrich) and Histopaque-1119 (density: 1.119 g/mL, Sigma-Aldrich) in a 1:1 ratio^37^. The purity of isolated neutrophils was checked via flow cytometry analyses with Ly-6G and CD11b antibodies (Supplementary Fig.3A). Isolated neutrophils from T1D and T2D were treated with 100ng/ml PMA (Sigma Aldrich, cat.no P1585) or 4μM Calcium Ionophore (CI) (Sigma Aldrich, cat.no C7522) for 2.5hrs (37°C, 5% CO_2_), in order to induce NETosis.

### Isolation of neutrophils from human peripheral blood

Peripheral blood from T1D and T2D patients and healthy donors was collected. Neutrophils were isolated using Histopaque-1119 (Sigma) and Percoll Plus (Sigma: P1644) density gradient methods. Blood was layered on Histopaque-1119 in a 1:1 ratio and centrifuged at 800 x g for 20 mins. The white blood cells were collected and washed with 1X PBS. Then the cells were layered on a Percoll gradient with concentrations 85, 80, 75, 70, and 65% and centrifuged for 20 minutes at 800 x g. Neutrophils were collected from Percoll layers positioned between 70-80% and cultured in RPMI 1640 with 10% FBS and 1% Pen/Strep^38^. The assays were repeated with different donors to obtain experimental replicates. Information related to T1D, T2D patients, and healthy controls is listed in the Supplementary Table 1.

### Inhibitors

For inhibitor experiments, cells were preincubated with pan-Pad inhibitor, CI-Amidine (200μM) (Sigma-506282), NOX inhibitor DPI (20μM) (Sigma-D2926), and mitochondrial superoxide scavenger MitoTempo (100μM) (Sigma- SML0737) for 1 hour before the induction of NETosis^39,40^.

### NETosis assay

SYTOX Green, a cell-impermeant dye (Thermo Fisher Scientific, cat. no. S7020), was used to measure NETosis in neutrophils derived from both T1D and T2D mice and in patient samples, with respective controls. Briefly, neutrophils were resuspended in RPMI 1640 with 10% FBS, 1% Pen/Strep. 2×10^4^ neutrophils were seeded per well in a 96-well plate (F-Bottom black, Fluotrac plate, Greiner bio-one) and SYTOX Green was added at a final concentration of 2 µM. Subsequently, NETosis was induced by 100 ng/ml PMA (Sigma-Aldrich, cat. no. P1585) or 4 μM Calcium Ionophore (Sigma-Aldrich, cat. no. C7522) for 2.5 hours^41^. DMSO was used as vehicle. NETs were quantified by measuring fluorescence using POLARstar OMEGA Fluorescent plate reader.

### Flow cytometry analysis

Neutrophils purity was checked by the FACS analyses using CD11b and Ly-6G (Supplementary Fig. 3A). Briefly, neutrophils were incubated with a mouse Fc-receptor blocker (Miltenyi Biotec) for 30 minutes at 4°C and stained with fluorescently conjugated antibody against CD11b (FITC-conjugated anti-CD11b, CST) and Ly-6G (PE anti-mouse Ly-6G Antibody, BioLegend) for 30 mins at 4°C^42^. Cells were washed twice with FACS buffer (1× PBS, supplemented with 2% FBS). Samples were analyzed using the BD FACSAria^TM^ cell sorter. Data were analyzed using FlowJo software.

### RNA isolation and gene expression analysis

Total RNA was isolated from neutrophils of T1D and T2D mice and from human patients using the RNeasy Mini Kit (Qiagen, Germany). RNA was reverse-transcribed using the iScript cDNA Synthesis Kit (Bio-Rad). RT–qPCR was performed using iTaq universal SYBR Green (BioRad) in a CFX Opus qPCR machine (Bio-Rad). The relative expression level of target genes was calculated using the 2^-ΔΔCt method^43^. 18S rRNA was used as the housekeeping gene control. All primers were synthesized by IDT (primer list). The list of primers used in this study is listed in Supplementary Table 2.

### Immunofluorescence analysis

1×10^5^ neutrophils were seeded onto a sterile coverslip in a 24-well plate. Further, NETosis was induced by incubating the cells with 100 ng/ml PMA or 4 μM CI for 2.5 hrs. Cells were fixed with 4% paraformaldehyde (Himedia: GRM3660-50G) for 4 hours at room temperature. Cells were quenched with 10 mM Tris-Cl and permeabilized with 0.1% Triton X-100. Cells were stained with primary antibodies against PADI4/PAD4 (abcam: ab96754, 1:250), citrullinated histone H3 (abcam: ab281584, 1:2000), myeloperoxidase (abcam: ab208670, 1:100), Ly-6G (Biolegend: 127601,1:400) for 30 minutes at 37°C. Cells were washed twice with PBST (0.1% Triton X-100 in PBS) for 10 minutes. Coverslips were incubated with secondary antibodies against Goat anti-Mouse Alexa Fluor™ 568 (Invitrogen: A-11019, 1:480), anti-Rabbit IgG Alexa Fluor™ 488 (Invitrogen: A-11070, 1:480) for 1.5 hours at 37°C. DNA was stained with DAPI (Thermo Fisher Scientific:62248, 1:1000). The coverslip was mounted with Fluoromount (Sigma: F4680). Images were taken using a LSM 980 Airyscan 2 (Zeiss, Germany). Images were analyzed with ImageJ software^20,38,44^.

### Western Blot analysis

2×10^6^ neutrophils were seeded in a 6-well plate in RPMI 1640 media containing 10% FBS, 1% Pen/Strep. Cells were stimulated with 100 ng/mL PMA or 4 μM CI for 2.5 hours at 37°C to induce NETosis. The cells were collected and washed with ice-cold 1X PBS. Cells were lysed with NP-40 lysis buffer (150mM NaCl, 1% NP-40, 50mM Tris-Cl, pH 8) containing complete protease inhibitor mixture (Roche: 5056489001) supplemented with NaVO3 (1 mM) (Sigma: 567540), leupeptin (5 μg/ml) (Sigma: L2884), pepstatin A (1 μg/ml) (Sigma: PEPS-RO), PMSF (1 mM), β-Glycerophosphate disodium salt hydrate (2mM) (Sigma: G9422) on ice for 30 mins. Lysates were boiled at 95 °C for 10 minutes. Equal amount of protein lysates was resolved on 10% SDS-PAGE gel and electroblotted on PVDF membranes. The membranes were incubated with primary antibodies against PADI4/PAD4 (1:5000), citrullinated histone H3 (1:2000), Myeloperoxidase (1:1000), GAPDH (CST:2118S, 1:3000) overnight at 4°C. The blots were washed twice with PBST and then incubated with HRP-conjugated secondary antibodies against rabbit IgG (CST:7074, 1:5000). The blots were developed with enhanced chemiluminescence substrate (BIO-RAD, 1705061) using Chemidoc (BioRad). Blots were quantified using ImageJ software^13^.

### Detection of ROS Production

In order to detect total ROS production and mitochondrial ROS production, isolated neutrophils were preincubated with 25μM Dihydrorhodamine 123 (DHR123) (Sigma: D1054) and 1μM MitoSOX Mitochondrial Superoxide Indicator (Thermo Fisher Scientific: M36007), respectively, for 30 minutes. The cells were activated with 100 ng/ml PMA or 4 μM CI for 2.5 hrs. Confocal imaging was done for qualitative analysis, and plate reader assays were performed for quantitative analysis as described before^13^.

### STZ-induced Pad4^-/-^ mice generation

T1D was induced in Pad4^-/-^ mice by five consecutive low doses of STZ injections. Briefly, 10-week-old male Pad4^-/-^ mice were fasted for 6 hours and then injected intraperitoneally with either vehicle or STZ (50 mg/kg/day) for five consecutive days. Fasting blood glucose levels and body weight were monitored throughout the study. Mice with fasting blood glucose levels above 300 mg/dL were considered diabetic and used for further experiments.

### Statistical analyses

Based on our prior experience, we used four to six mice per group for the *in vivo* and *ex vivo* experiments. All *ex vivo* experiments were performed at least three times unless indicated otherwise. Statistical analyses and plots were generated using GraphPad Prism 10.0 software (GraphPad Prism Software Inc., San Diego, CA) unless stated otherwise. The normality of each sample group was assessed using the Shapiro–Wilk normality test before comparing groups. For statistical comparison between two groups, unpaired Student’s t-test (two tails) was used to determine the significance of intergroup differences. For comparisons of three or more groups, when variances were similar among the groups, multiple groups were analyzed by one-way analysis of variance (ANOVA), followed by Tukey’s or Dunnett’s post hoc tests. The results are expressed as the means ± SD, and *p*-values < 0.05 were considered statistically significant for all tests used.

## Supporting information

supplementary merged

## Acknowledgements

This study was supported by grants from ANRF (ANRF/IRG/2024/000321/LS) and, in part, by a grant from DBT/Wellcome Trust India Alliance (IA/I/19/1/50427), SERB (SRG/2022/001063), DBT (BT/PR58221/BMS/85/1402/2025), and IISER-Mohali Start-up (to SD). PS is supported by DBT-Ramalingaswami Re-entry fellowship and BRIC-RGCB intramural funding. SG, AP, SD (1), and DS are supported by UGC and CSIR, Government of India. SD (2) is supported by IISER-Mohali, KK is supported by India Alliance. SD (3) is a DBT/Wellcome Trust India Alliance Intermediate Fellow. We acknowledge support provided by the Animal House, microscopy (DBT-FIST), and FACS facilities at IISER-Mohali.

## Author contributions

SG designed and performed most of the experiments, the data analyses, and wrote the manuscript. AP, DS, SD(1), SD(2), KK, and PK performed experiments and analyzed the data. AM and SB provided the patient samples and edited the manuscript. PS analyzed, wrote, and edited the manuscript. SD (3) conceptualized the work, wrote and edited the manuscript, acquired funding, and supervised the study. SD (3) is the guarantor of this work and, as such, had full access to all the data in the study and takes responsibility for the integrity of the data and the accuracy of the data analysis.

## References

1. American Diabetes A. 2. Classification and Diagnosis of Diabetes: Standards of Medical Care in Diabetes-2018. Diabetes Care. 2018;41:S13–S27. doi: 10.2337/dc18-S002

2. Atkinson MA, Eisenbarth GS, Michels AW. Type 1 diabetes. Lancet. 2014;383:69–82. doi: 10.1016/S0140-6736(13)60591-7

3. Fuchs TA, Abed U, Goosmann C, Hurwitz R, Schulze I, Wahn V, Weinrauch Y, Brinkmann V, Zychlinsky A. Novel cell death program leads to neutrophil extracellular traps. J Cell Biol. 2007;176:231–241. doi: 10.1083/jcb.200606027

4. Brinkmann V, Reichard U, Goosmann C, Fauler B, Uhlemann Y, Weiss DS, Weinrauch Y, Zychlinsky A. Neutrophil extracellular traps kill bacteria. Science. 2004;303:1532–1535. doi: 10.1126/science.1092385

5. Yipp BG, Kubes P. NETosis: how vital is it? Blood. 2013;122:2784–2794. doi: 10.1182/blood-2013-04-457671

6. Franck G, Mawson TL, Folco EJ, Molinaro R, Ruvkun V, Engelbertsen D, Liu X, Tesmenitsky Y, Shvartz E, Sukhova GK, et al. Roles of PAD4 and NETosis in Experimental Atherosclerosis and Arterial Injury: Implications for Superficial Erosion. Circ Res. 2018;123:33–42. doi: 10.1161/CIRCRESAHA.117.312494

7. Munks MW, McKee AS, Macleod MK, Powell RL, Degen JL, Reisdorph NA, Kappler JW, Marrack P. Aluminum adjuvants elicit fibrin-dependent extracellular traps in vivo. Blood. 2010;116:5191–5199. doi: 10.1182/blood-2010-03-275529

8. Wong SL, Demers M, Martinod K, Gallant M, Wang Y, Goldfine AB, Kahn CR, Wagner DD. Diabetes primes neutrophils to undergo NETosis, which impairs wound healing. Nat Med. 2015;21:815–819. doi: 10.1038/nm.3887

9. Fadini GP, Menegazzo L, Rigato M, Scattolini V, Poncina N, Bruttocao A, Ciciliot S, Mammano F, Ciubotaru CD, Brocco E, et al. NETosis Delays Diabetic Wound Healing in Mice and Humans. Diabetes. 2016;65:1061–1071. doi: 10.2337/db15-0863

10. Liu C, Yalavarthi S, Tambralli A, Zeng L, Rysenga CE, Alizadeh N, Hudgins L, Liang W, NaveenKumar SK, Shi H, et al. Inhibition of neutrophil extracellular trap formation alleviates vascular dysfunction in type 1 diabetic mice. Sci Adv. 2023;9:eadj1019. doi: 10.1126/sciadv.adj1019

11. Karima M, Kantarci A, Ohira T, Hasturk H, Jones VL, Nam BH, Malabanan A, Trackman PC, Badwey JA, Van Dyke TE. Enhanced superoxide release and elevated protein kinase C activity in neutrophils from diabetic patients: association with periodontitis. J Leukoc Biol. 2005;78:862–870. doi: 10.1189/jlb.1004583

12. Eizirik DL, Colli ML, Ortis F. The role of inflammation in insulitis and beta-cell loss in type 1 diabetes. Nat Rev Endocrinol. 2009;5:219–226. doi: 10.1038/nrendo.2009.21

13. Douda DN, Khan MA, Grasemann H, Palaniyar N. SK3 channel and mitochondrial ROS mediate NADPH oxidase-independent NETosis induced by calcium influx. Proc Natl Acad Sci U S A. 2015;112:2817–2822. doi: 10.1073/pnas.1414055112

14. Metzler KD, Goosmann C, Lubojemska A, Zychlinsky A, Papayannopoulos V. A myeloperoxidase-containing complex regulates neutrophil elastase release and actin dynamics during NETosis. Cell Rep. 2014;8:883–896. doi: 10.1016/j.celrep.2014.06.044

15. Wang Y, Li M, Stadler S, Correll S, Li P, Wang D, Hayama R, Leonelli L, Han H, Grigoryev SA, et al. Histone hypercitrullination mediates chromatin decondensation and neutrophil extracellular trap formation. J Cell Biol. 2009;184:205–213. doi: 10.1083/jcb.200806072

16. Arita K, Hashimoto H, Shimizu T, Nakashima K, Yamada M, Sato M. Structural basis for Ca(2+)-induced activation of human PAD4. Nat Struct Mol Biol. 2004;11:777–783. doi: 10.1038/nsmb799

17. Dwivedi N, Radic M. Citrullination of autoantigens implicates NETosis in the induction of autoimmunity. Ann Rheum Dis. 2014;73:483–491. doi: 10.1136/annrheumdis-2013-203844

18. Popp SK, Vecchio F, Brown DJ, Fukuda R, Suzuki Y, Takeda Y, Wakamatsu R, Sarma MA, Garrett J, Giovenzana A, et al. Circulating platelet-neutrophil aggregates characterize the development of type 1 diabetes in humans and NOD mice. JCI Insight. 2022;7. doi: 10.1172/jci.insight.153993

19. Lewis HD, Liddle J, Coote JE, Atkinson SJ, Barker MD, Bax BD, Bicker KL, Bingham RP, Campbell M, Chen YH, et al. Inhibition of PAD4 activity is sufficient to disrupt mouse and human NET formation. Nat Chem Biol. 2015;11:189–191. doi: 10.1038/nchembio.1735

20. Neubert E, Meyer D, Rocca F, Gunay G, Kwaczala-Tessmann A, Grandke J, Senger-Sander S, Geisler C, Egner A, Schon MP, et al. Chromatin swelling drives neutrophil extracellular trap release. Nat Commun. 2018;9:3767. doi: 10.1038/s41467-018-06263-5

21. Metzler KD, Fuchs TA, Nauseef WM, Reumaux D, Roesler J, Schulze I, Wahn V, Papayannopoulos V, Zychlinsky A. Myeloperoxidase is required for neutrophil extracellular trap formation: implications for innate immunity. Blood. 2011;117:953–959. doi: 10.1182/blood-2010-06-290171

22. Egesten A, Breton-Gorius J, Guichard J, Gullberg U, Olsson I. The heterogeneity of azurophil granules in neutrophil promyelocytes: immunogold localization of myeloperoxidase, cathepsin G, elastase, proteinase 3, and bactericidal/permeability increasing protein. Blood. 1994;83:2985–2994.

23. Papayannopoulos V, Metzler KD, Hakkim A, Zychlinsky A. Neutrophil elastase and myeloperoxidase regulate the formation of neutrophil extracellular traps. J Cell Biol. 2010;191:677–691. doi: 10.1083/jcb.201006052

24. Keshari RS, Verma A, Barthwal MK, Dikshit M. Reactive oxygen species-induced activation of ERK and p38 MAPK mediates PMA-induced NETs release from human neutrophils. J Cell Biochem. 2013;114:532–540. doi: 10.1002/jcb.24391

25. Furman BL. Streptozotocin-Induced Diabetic Models in Mice and Rats. Curr Protoc Pharmacol. 2015;70:5 47 41–45 47 20. doi: 10.1002/0471141755.ph0547s70

26. Festa A, D’Agostino R, Jr., Howard G, Mykkanen L, Tracy RP, Haffner SM. Chronic subclinical inflammation as part of the insulin resistance syndrome: the Insulin Resistance Atherosclerosis Study (IRAS). Circulation. 2000;102:42–47. doi: 10.1161/01.cir.102.1.42

27. Butler AE, Janson J, Bonner-Weir S, Ritzel R, Rizza RA, Butler PC. Beta-cell deficit and increased beta-cell apoptosis in humans with type 2 diabetes. Diabetes. 2003;52:102–110. doi: 10.2337/diabetes.52.1.102

28. Brownlee M. Biochemistry and molecular cell biology of diabetic complications. Nature. 2001;414:813–820. doi: 10.1038/414813a

29. Hotamisligil GS, Shargill NS, Spiegelman BM. Adipose expression of tumor necrosis factor-alpha: direct role in obesity-linked insulin resistance. Science. 1993;259:87–91. doi: 10.1126/science.7678183

30. Lee KH, Kronbichler A, Park DD, Park Y, Moon H, Kim H, Choi JH, Choi Y, Shim S, Lyu IS, et al. Neutrophil extracellular traps (NETs) in autoimmune diseases: A comprehensive review. Autoimmun Rev. 2017;16:1160–1173. doi: 10.1016/j.autrev.2017.09.012

31. Papayannopoulos V. Neutrophil extracellular traps in immunity and disease. Nat Rev Immunol. 2018;18:134–147. doi: 10.1038/nri.2017.105

32. Jorch SK, Kubes P. An emerging role for neutrophil extracellular traps in noninfectious disease. Nat Med. 2017;23:279–287. doi: 10.1038/nm.4294

33. Neeli I, Khan SN, Radic M. Histone deimination as a response to inflammatory stimuli in neutrophils. J Immunol. 2008;180:1895–1902. doi: 10.4049/jimmunol.180.3.1895

34. Neeli I, Radic M. Opposition between PKC isoforms regulates histone deimination and neutrophil extracellular chromatin release. Front Immunol. 2013;4:38. doi: 10.3389/fimmu.2013.00038

35. Kenny EF, Herzig A, Kruger R, Muth A, Mondal S, Thompson PR, Brinkmann V, Bernuth HV, Zychlinsky A. Diverse stimuli engage different neutrophil extracellular trap pathways. Elife. 2017;6. doi: 10.7554/eLife.24437

36. Lei L, Liu J, Li Y, Huang B, Zou Z. Neutrophils and neutrophil extracellular traps in diabetes mellitus and its complications: Mechanisms and therapeutic implications. iScience. 2026;29:115585. doi: 10.1016/j.isci.2026.115585

37. Zhao X, Yang L, Chang N, Hou L, Zhou X, Yang L, Li L. Neutrophils undergo switch of apoptosis to NETosis during murine fatty liver injury via S1P receptor 2 signaling. Cell Death Dis. 2020;11:379. doi: 10.1038/s41419-020-2582-1

38. Thiam HR, Wong SL, Qiu R, Kittisopikul M, Vahabikashi A, Goldman AE, Goldman RD, Wagner DD, Waterman CM. NETosis proceeds by cytoskeleton and endomembrane disassembly and PAD4-mediated chromatin decondensation and nuclear envelope rupture. Proc Natl Acad Sci U S A. 2020;117:7326–7337. doi: 10.1073/pnas.1909546117

39. Bork F, Greve CL, Youn C, Chen S, V NCL, Wang Y, Fischer B, Nasri M, Focken J, Scheurer J, et al. naRNA-LL37 composite DAMPs define sterile NETs as self-propagating drivers of inflammation. EMBO Rep. 2024;25:2914–2949. doi: 10.1038/s44319-024-00150-5

40. Lefrancais E, Mallavia B, Zhuo H, Calfee CS, Looney MR. Maladaptive role of neutrophil extracellular traps in pathogen-induced lung injury. JCI Insight. 2018;3. doi: 10.1172/jci.insight.98178

41. Azzouz D, Palaniyar N. ROS and DNA repair in spontaneous versus agonist-induced NETosis: Context matters. Front Immunol. 2022;13:1033815. doi: 10.3389/fimmu.2022.1033815

42. Zhu YP, Speir M, Tan Z, Lee JC, Nowell CJ, Chen AA, Amatullah H, Salinger AJ, Huang CJ, Wu G, et al. NET formation is a default epigenetic program controlled by PAD4 in apoptotic neutrophils. Sci Adv. 2023;9:eadj1397. doi: 10.1126/sciadv.adj1397

43. Das S, Zhang E, Senapati P, Amaram V, Reddy MA, Stapleton K, Leung A, Lanting L, Wang M, Chen Z, et al. A Novel Angiotensin II-Induced Long Noncoding RNA Giver Regulates Oxidative Stress, Inflammation, and Proliferation in Vascular Smooth Muscle Cells. Circ Res. 2018;123:1298–1312. doi: 10.1161/CIRCRESAHA.118.313207

44. Munir S, Basu A, Maity P, Krug L, Haas P, Jiang D, Strauss G, Wlaschek M, Geiger H, Singh K, et al. TLR4-dependent shaping of the wound site by MSCs accelerates wound healing. EMBO Rep. 2020;21:e48777. doi: 10.15252/embr.201948777

