## supplementary merged for "PAD4-mediated histone citrullination contributes to enhanced NETosis in type 2 diabetes but not in type 1 diabetes"

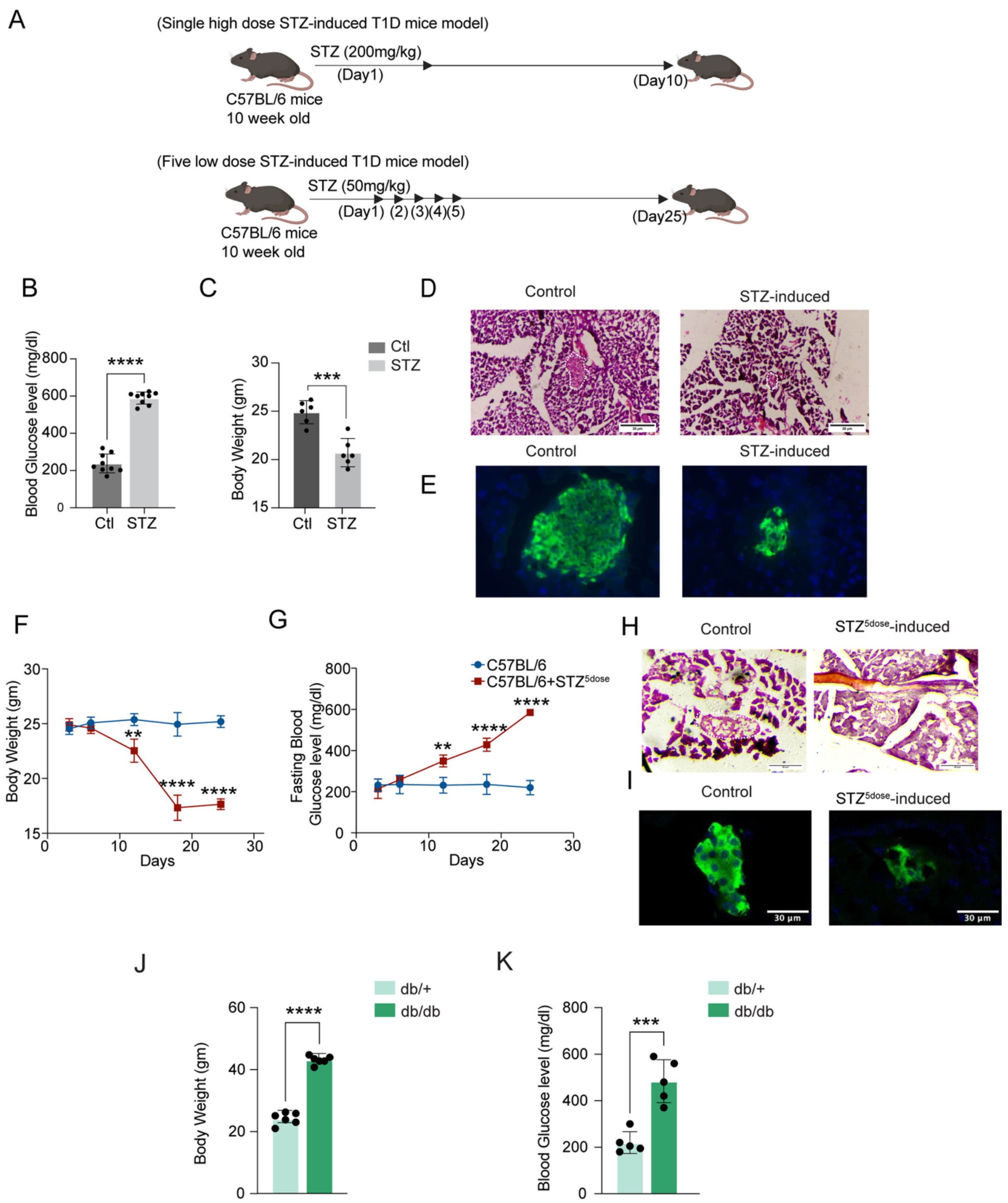


**Supplementary Fig. 1. Characterization of STZ-induced diabetes in C57BL/6 mice.** (A) Experimental strategy of single high dose STZ (200mg/kg) and five low dose STZ (50mg/kg) induced T1D mice model. (B) Diabetes was defined as high blood glucose >300 mg/dl, single high dose STZ-treated mice became diabetic on 10$^{th}$ day after treatment (n=9). (C) Body weight was significantly reduced on 10$^{th}$ day after STZ treatment (n=6). (D) H&E staining showing reduction in pancreatic islets cells upon

STZ induction. (E) Representative IF images showing a marked reduction of insulin-producing β cells and disrupted islet morphology in the pancreas of STZ-treated mice. (F and G) Decrease in body weight and increase in fasting blood glucose in 5 low dose STZ-treated C57BL/6 mice (n=3). (H) H&E staining showing reduction in pancreatic islets cells upon 5 lose dose STZ induction. (I) Representative IF images showing a marked reduction of insulin-producing β cells and disrupted islet morphology in the pancreas of 5 low dose STZ-treated mice. (J) Body weight and (K) Blood glucose were significantly higher in 8-10 weeks old db/db male mice (n=6). *$P$ < 0.05, **$P$ < 0.01, ***$P$ < 0.001, ****$P$< 0.0001. Data represented as means ± SD.

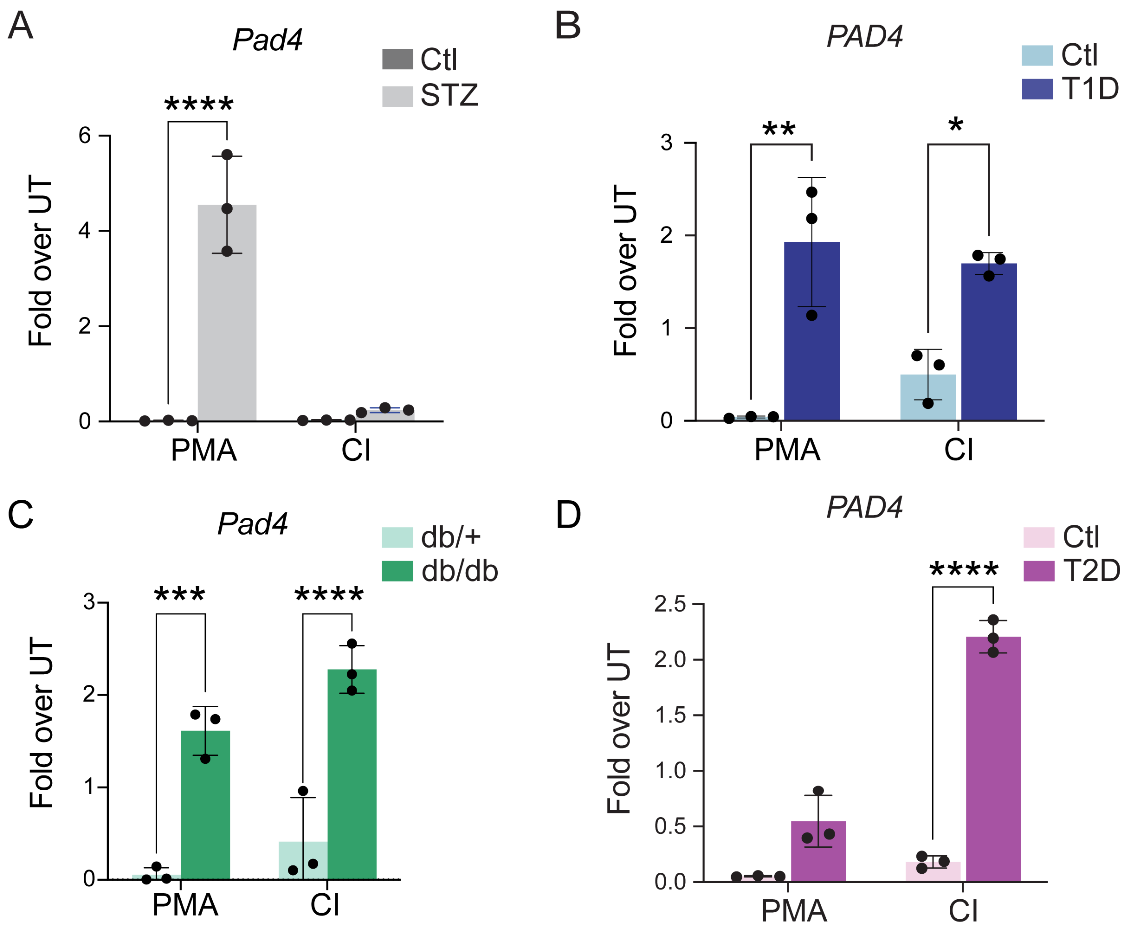


**Supplementary Fig. 2.** (A and C) RT-qPCR data showing *Pad4* expression in neutrophils from STZ-induced and db/db mice respectively (n=3). (B and D) *PAD4* expression in neutrophils from T1D and T2D patient blood (n=3). *$P$ < 0.05, **$P$ < 0.01, ***$P$ < 0.001, ****$P$< 0.0001. Data represented as means ± SD.

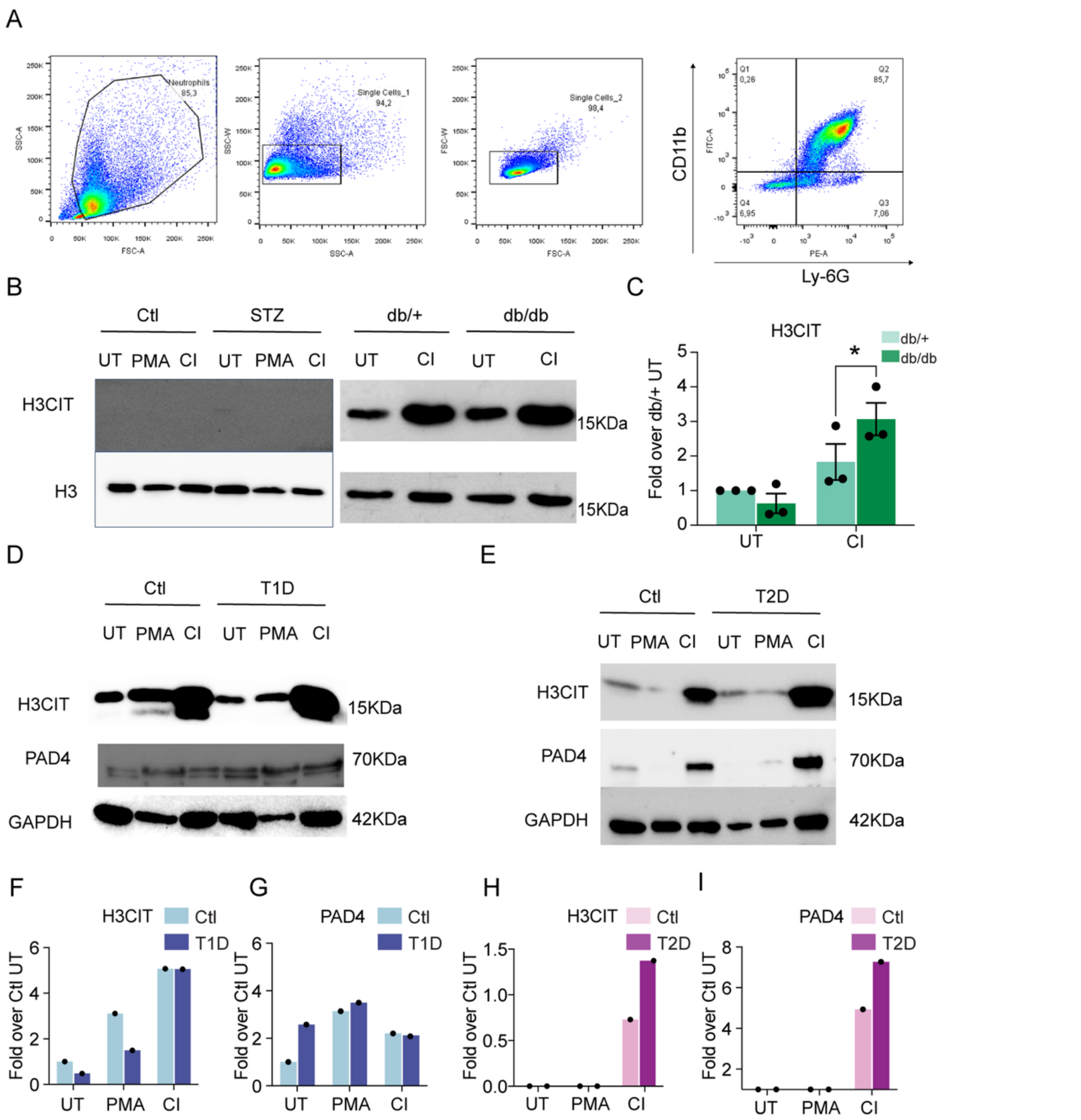


**Supplementary Fig. 3.** (A) Representative FACS plots of bone marrow derived neutrophils. CD11b/Ly-6G show >80 % neutrophil purity. (B and C) Western blot analysis of H3CIT expression in neutrophils from STZ-induced and db/db mice. (D and E) Western blot analysis of H3CIT and PAD4 expression in neutrophils from healthy individuals and T1D patients and T2D patients respectively. For each blot neutrophils are isolated and pooled from 4 patients. (F-I) quantification of (D and E). *$P$ < 0.05, **$P$ < 0.01, ***$P$ < 0.001, ****$P$< 0.0001. Data represented as means ± SD.

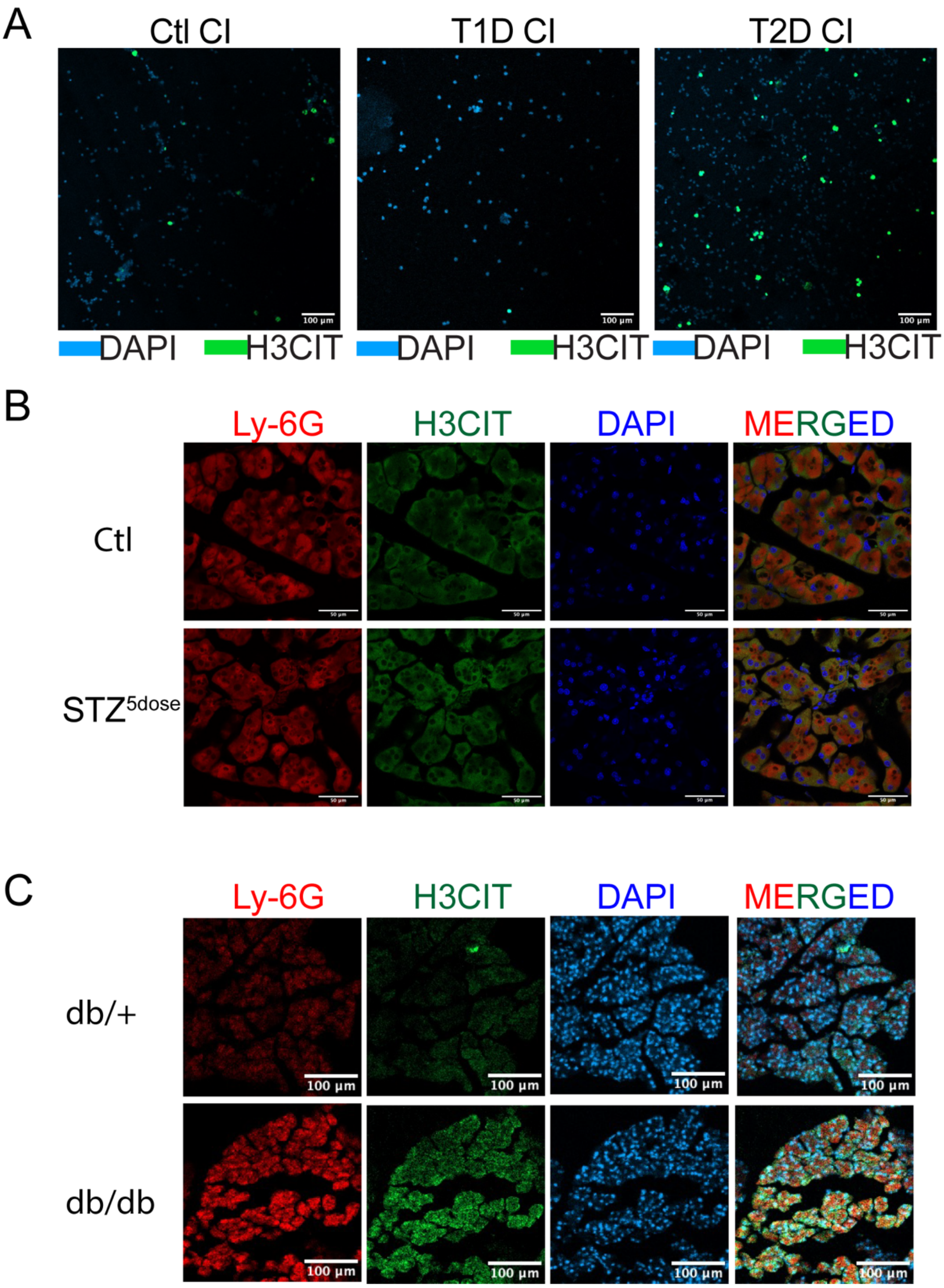


**Supplementary Fig. 4**. (A) Representative IF images of neutrophils isolated from healthy individual, T1D patient and T2D patient and stimulated with CI, showing histone 3 citrullination positive cells, scale bar, 100 μm. (B) Representative IF image showing H3CIT abundance in pancreatic sections isolated from STZ-treated mice. (C) IF image showing H3CIT abundance in pancreatic sections isolated db/+ and db/db mice, scale bar, 100 μm.

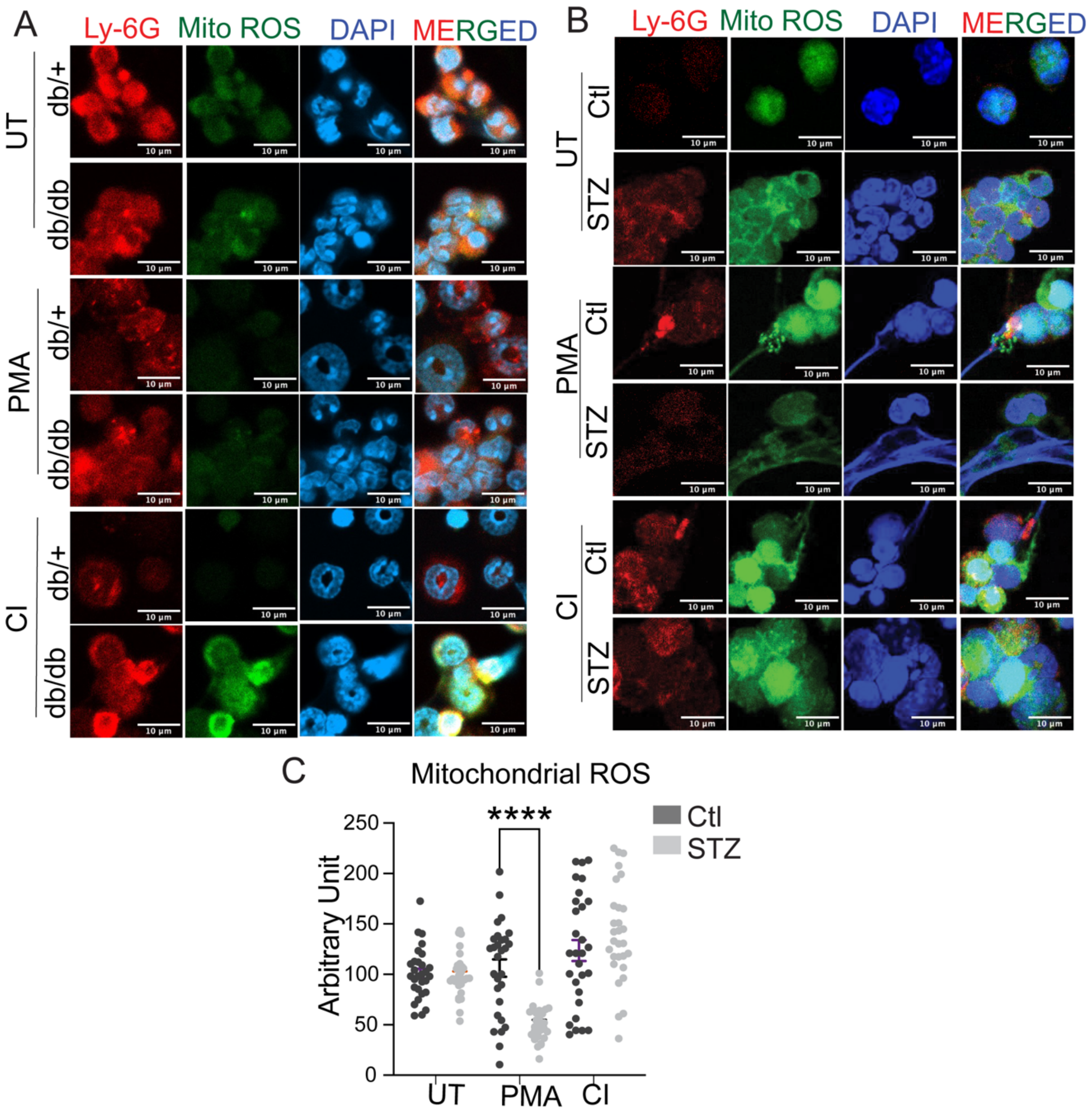


**Supplementary Fig. 5.** (A) IF showing mitochondrial ROS expression in neutrophils isolated from db/db mice. (B) IF showing mitochondrial ROS expression in neutrophils isolated from STZ-induced mice and (C) its quantification (n =30 cells), scale bar, 10 µm. **P* < 0.05, ***P* < 0.01, ****P* < 0.001, *****P*< 0.0001. Data represented as means ± SD.

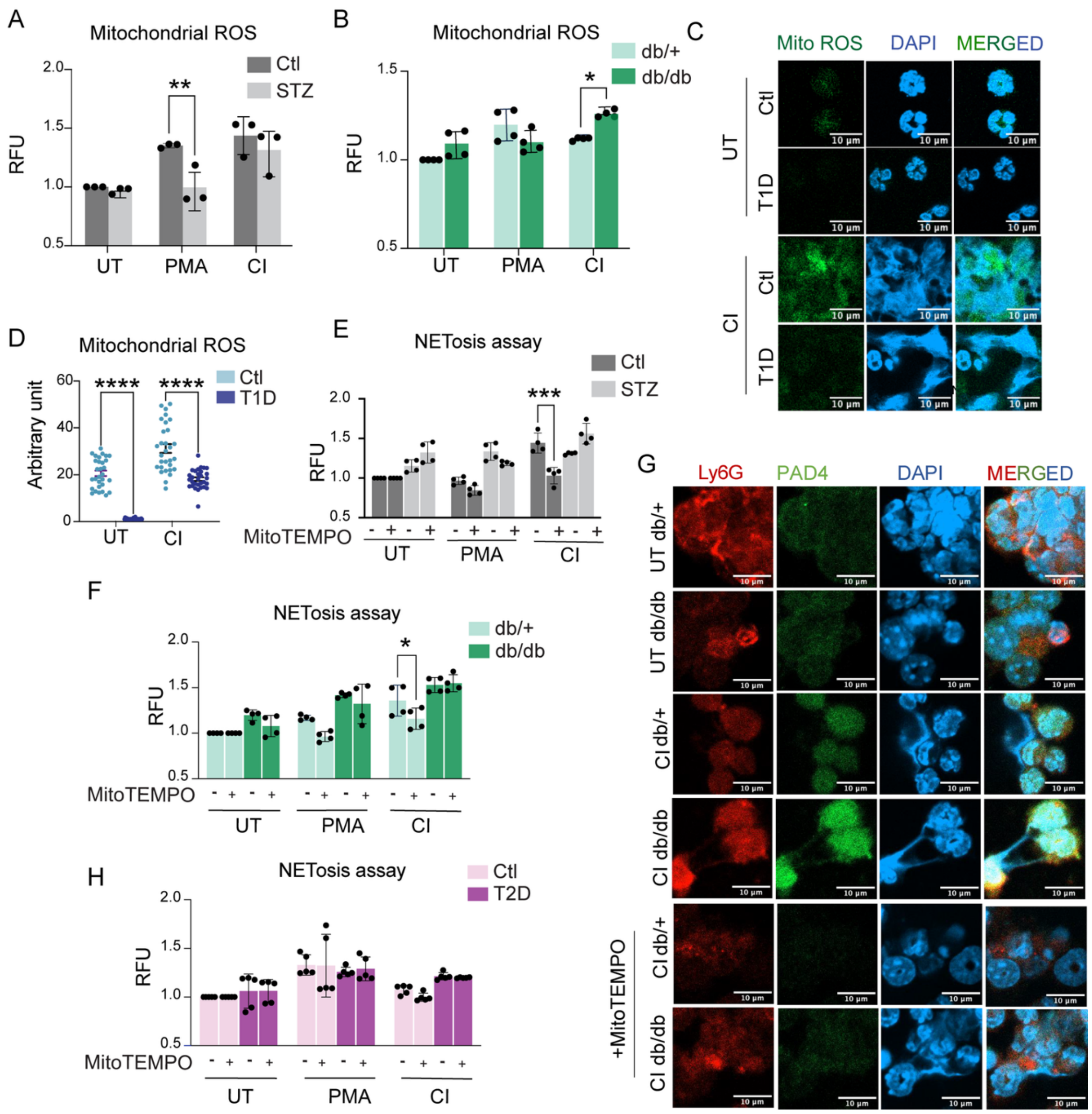


**Supplementary Fig. 6**. (A and B) Mitochondrial ROS levels in PMA- and CI-treated neutrophils form STZ-induced and db/db mice respectively measured by MitoSOX dye plate reader assay, (n=3-4). (C) Fluorescence images showing mitochondrial ROS (mtROS) level in CI-treated neutrophils from T1D patients and (D) its quantification (n=30 cells). (E and F) SYTOX assay showing role of Mitochondrial ROS in PMA- and CI-treated neutrophils with mtROS inhibitor MitoTEMPO in NET formation in STZ-induced and db/db mice (n=4). (G) IF showing reduced PAD4 expression on mtROS inhibitor (MitoTEMPO) in neutrophils from db/db mice. (H) SYTOX assay showing role of Mitochondrial ROS upon treatment of neutrophils with mtROS inhibitor MitoTEMPO in NET formation in T2D patients (n=5). *$P$ < 0.05, **$P$ < 0.01, ***$P$ < 0.001, ****$P$< 0.0001). Data represented as means ± SD.

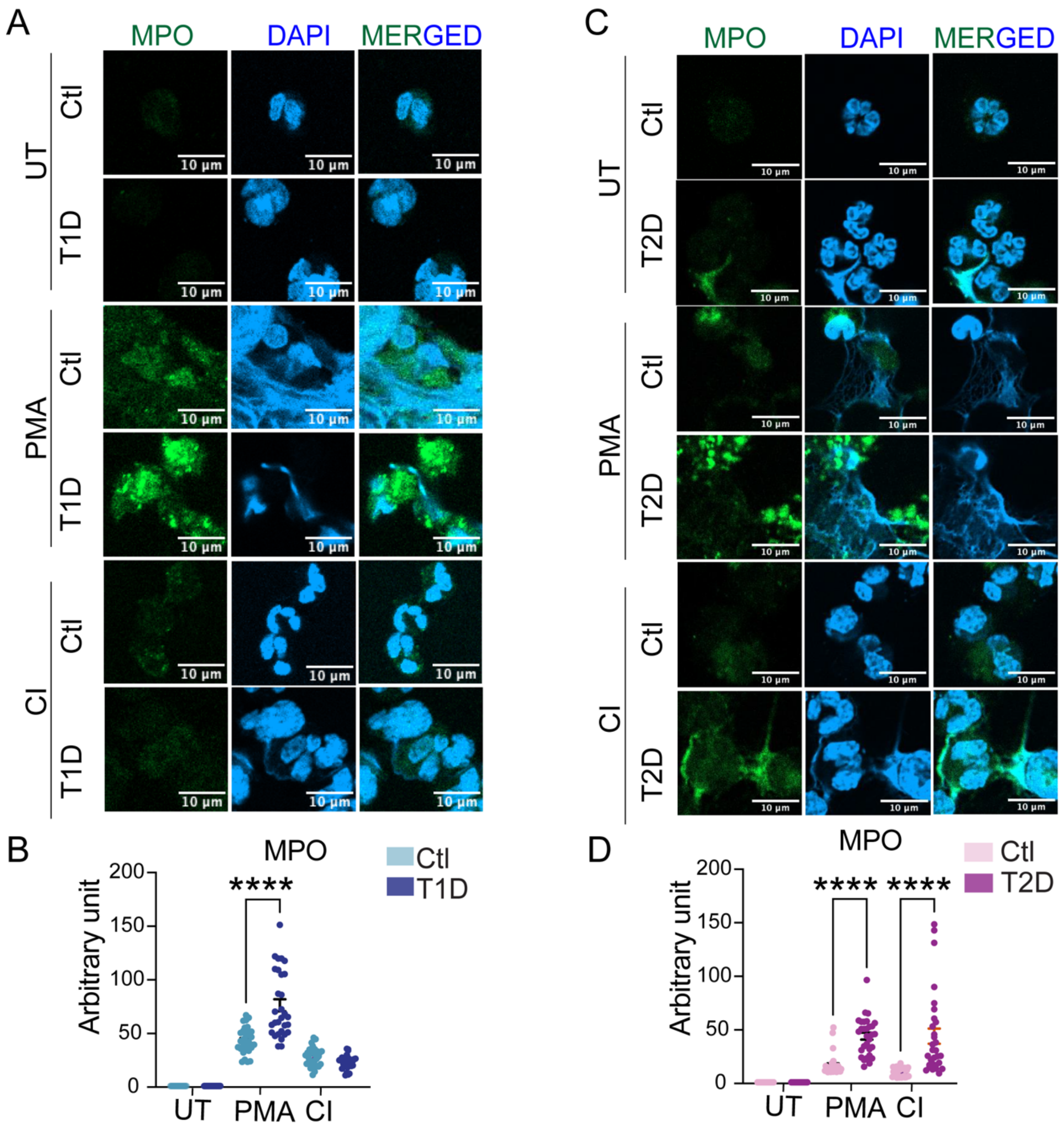


**Supplementary Fig. 7.** (A) IF showing MPO expression in neutrophils isolated from Type I diabetic patients and (B) its quantification (n =30 cells), scale bar, 10 µm. (C) IF showing MPO expression in neutrophils isolated from Type II diabetic patients and (D) its quantification (n =30 cells), scale bar, 10µm. **P* < 0.05, ***P* < 0.01, ****P* < 0.001, *****P*< 0.0001. Data represented as means ± SD.

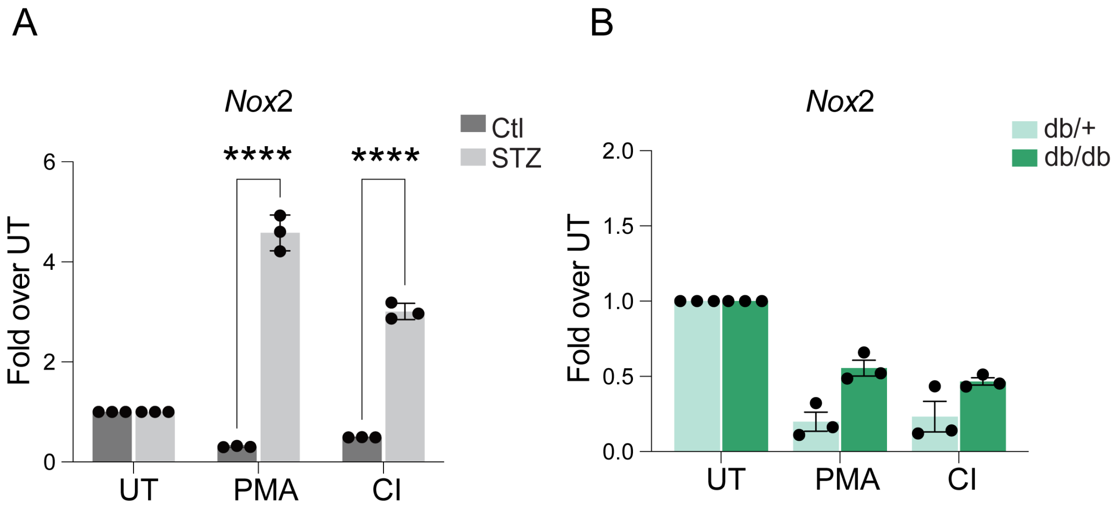


**Supplementary Fig. 8.** (A and B) RT-qPCR data showing *Nox*2 expression in neutrophils from STZ-induced and db/db mice respectively (n=3). *$P$ < 0.05, **$P$ < 0.01, ***$P$ < 0.001, ****$P$< 0.0001. Data represented as means ± SD.

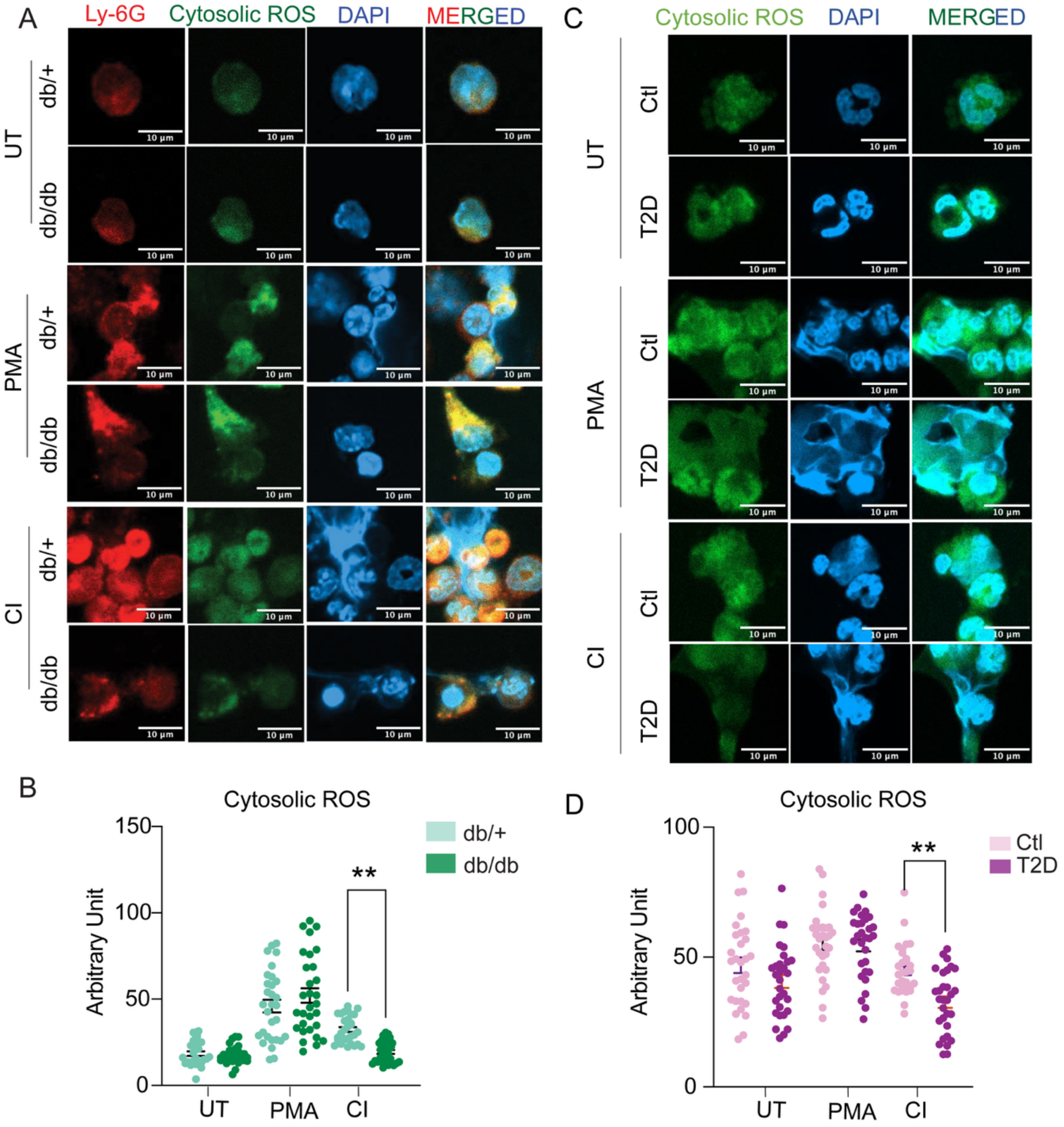


**Supplementary Fig. 9.** (A) IF showing cytosolic ROS expression in neutrophils isolated from db/db mice and (B) its quantification (n =30 cells), scale bar, 10 µm. (C) Fluorescence images showing cytosolic ROS expression in T2D patient sample and (D) its quantification (n =30 cells), scale bar, 10 µm. *$P < 0.05$, **$P < 0.01$, ***$P < 0.001$, ****$P< 0.0001$. Data represented as means ± SD.

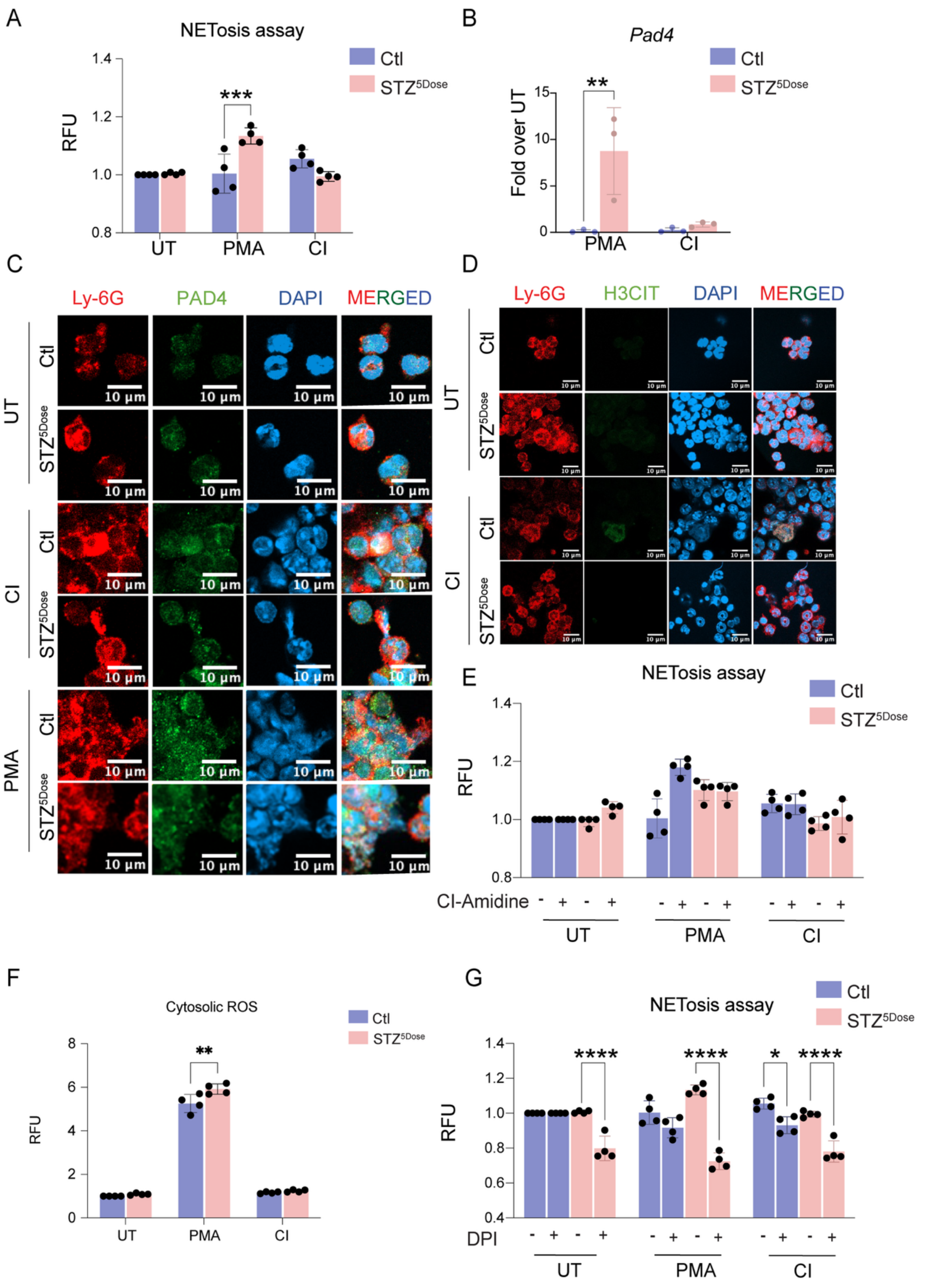


**Supplementary Fig. 10.** (A) SYTOX assay showing NET production in PMA- and CI-stimulated neutrophils isolated from 5 low dose STZ induced C57BL/6 mice (n=4). (B) RT-qPCR data showing *Pad4* expression in neutrophils from 5 low dose STZ-induced mice (n=3). (C) IF showing PAD4 expression in neutrophils isolated from 5 low dose STZ-induced, scale bar, 10 μm. (D) IF showing H3CIT expression in neutrophils isolated from 5 low dose STZ-induced mice, scale bar, 10 µm. (E) SYTOX assay showing effect of Pan-PAD4 inhibitor (Cl-Amidine) in NET formation in neutrophils from 5 low dose STZ-induced mice (n=4). (F) DHR123 (Dihydrorhodamine 123) expression showing cytosolic ROS levels in PMA- and Cl-treated neutrophils form 5 low dose STZ-induced mice (n=4). (G) SYTOX assay showing role of ROS in PMA- and Cl-treated neutrophils with ROS inhibitor DPI in NET formation in neutrophils of 5 low dose STZ-induced mice (n=4). *$P < 0.05$, **$P < 0.01$, ***$P < 0.001$, ****$P < 0.0001$. Data represented as means ± SD.

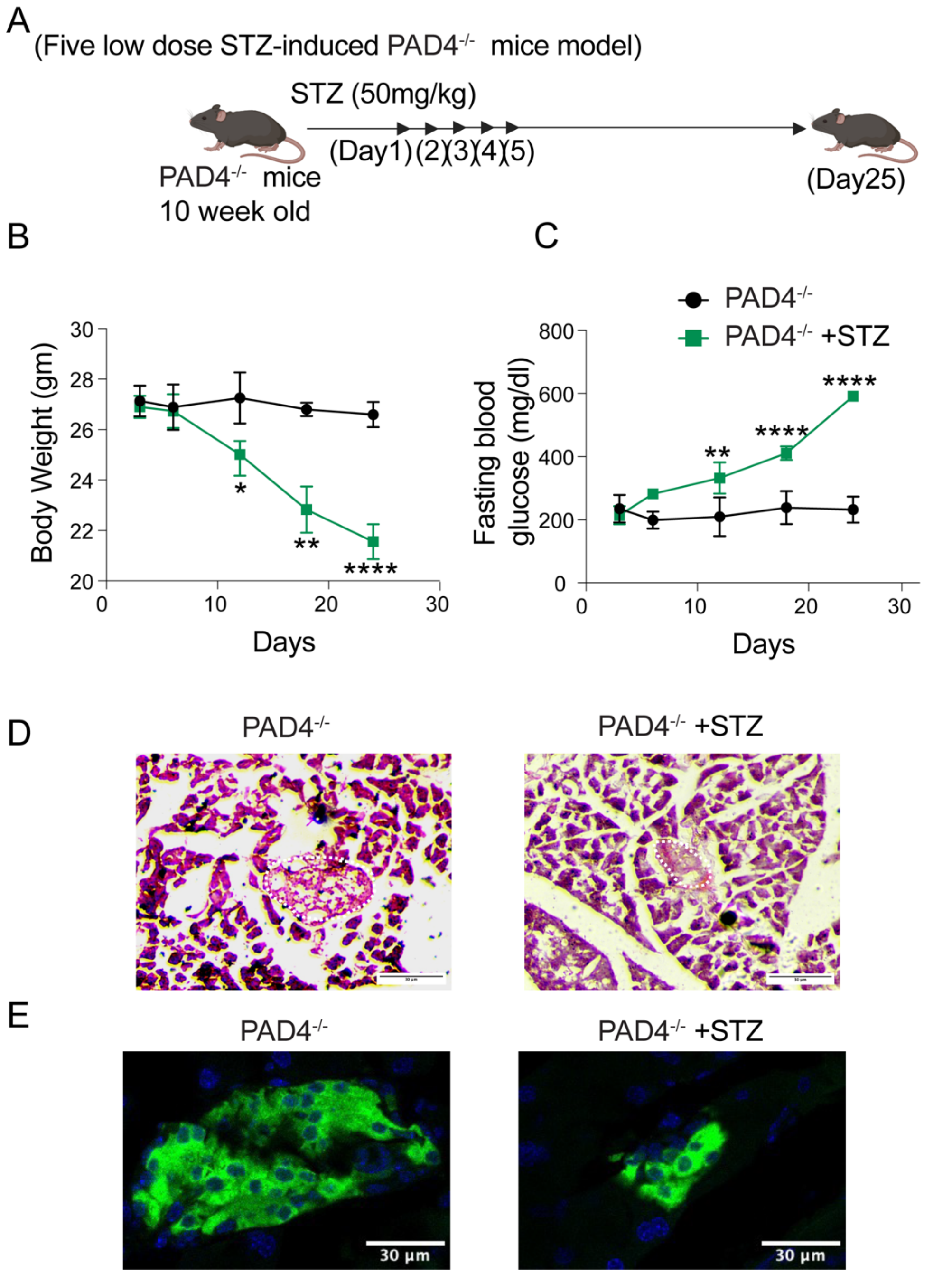


**Supplementary Fig. 11.** Characterization of STZ-induced diabetes in PAD4-/- mice. (A) Experimental strategy of single high dose STZ (200mg/kg) and five low dose STZ (50mg/kg) induced PAD4$^{-/-}$ T1D mice model (B and C) Decrease in body weight and increase in fasting blood glucose in 5 low dose STZ-treated PAD4$^{-/-}$ mice (n=3). (D) H&E staining showing reduction in pancreatic islets cells upon 5 low dose STZ induction. (E) Representative IF images showing a marked reduction of insulin-producing β cells and disrupted islet morphology in the pancreas of 5 low dose STZ-treated PAD4$^{-/-}$ mice. $P < 0.05$, $^{**}P < 0.01$, $^{***}P < 0.001$, $^{****}P < 0.0001$. Data represented as means ± SD.

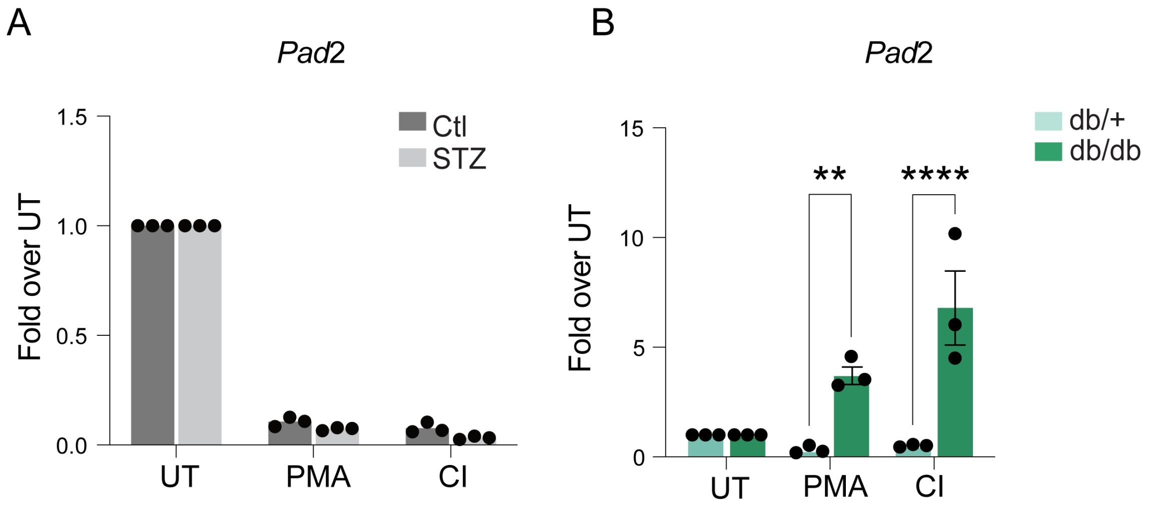


**Supplementary Fig. 12.** (A and B) RT-qPCR data showing *Pad2* expression in neutrophils from STZ-induced and db/db mice respectively (n=3). **P* < 0.05, ***P* < 0.01, ****P* < 0.001, *****P* < 0.0001. Data represented as means ± SD.

| S.No. | Donor | Age (years) | Gender | HbA1c levels (%) | Blood glucose fasting(mg/dl) | Disease duration (years) |
|---|---|---|---|---|---|---|
| 1 | T1D | 23 | F | 6.8 | 71 | 3 |
| 2 | T1D | 30 | F | 13.8 | 195 | 4 |
| 3 | T1D | 22 | M | 12.4 | 120 | 8 |
| 4 | T1D | 25 | F | 7.8 | 110 | 4 |
| 5 | T1D | 24 | F | 13.8 | 188 | 6 |
| 6 | T1D | 34 | F | 6.5 | 90 | 10 |
| 7 | T1D | 37 | M | 9.5 | 465 | 15 |
| 8 | T1D | 37 | M | 9 | 150 | 15 |
| 9 | T1D | 23 | F | 8.7 | 78 | 4 |
| 10 | T1D | 27 | F | 7 | 110 | 8 |
| 11 | T1D | 26 | F | 10.33 | 316 | 11 |
| 12 | T2D | 50 | M | 10.4 | 315.2 | 15 |
| 13 | T2D | 41 | F | 10.11 | 129 | 10 |
| 14 | T2D | 72 | F | 8.9 | 139 | 25 |
| 15 | T2D | 47 | F | 12.4 | 179 | 12 |
| 16 | T2D | 43 | F | 7.3 | 150 | 7 |
| 17 | T2D | 77 | M | 7.7 | 135 | 20 |
| 18 | T2D | 50 | F | 7.5 | 128 | 10 |
| 19 | T2D | 34 | M | 7.9 | 155 | 1 |
| 20 | T2D | 74 | M | 8.3 | 119 | 7 |
| 21 | T2D | 35 | M | 9.4 | 159 | 2 |
| 22 | T2D | 53 | F | 6.6 | 123 | 2 |
| 23 | T2D | 50 | F | 8.2 | 130 | 24 |
| 24 | T2D | 42 | F | 6.85 | 148 | 5 |
| 25 | T2D | 55 | M | 6.7 | 177 | 10 |
| 26 | T2D | 72 | F | 8.98 | 189 | 28 |
| 27 | Healthy | 25 | F | 5.5 | 88 | Nil |
| 28 | Healthy | 32 | F | 4.78 | 75 | Nil |
| 29 | Healthy | 35 | F | 5.8 | 94 | Nil |
| 30 | Healthy | 44 | M | 4.67 | 91 | Nil |
| 31 | Healthy | 24 | M | 5.5 | 101 | Nil |
| 32 | Healthy | 27 | F | 5.7 | 86 | Nil |
| 33 | Healthy | 50 | M | 5.9 | 107 | Nil |
| 34 | Healthy | 38 | M | 4.6 | 97 | Nil |
| 35 | Healthy | 29 | F | 4.7 | 82 | Nil |
| 36 | Healthy | 36 | M | 5.6 | 105 | Nil |
| 37 | Healthy | 34 | F | 5.47 | 98 | Nil |
| 38 | Healthy | 25 | F | 4.9 | 87 | Nil |
| 39 | Healthy | 29 | F | 5.1 | 110 | Nil |
| 40 | Healthy | 23 | M | 4.4 | 79 | Nil |

**Supplementary Table 1.** Characteristics of healthy and diabetic blood donors.

| | Healthy individuals | T2D individuals | T1D individuals |
|---|---|---|---|
| HbA1c(%) | <6.0 | >6.5 | >6.5 |
| Glucose fasting (mg/dl) | <110 | >119 | >119 |

**Supplementary Table 2.** Parameters for T1D and T2D patients

| Name | Sequence (5' to 3') |
|---|---|
| m*Pad4*-F | CGGAATGGACTTTGAGGATGAC |
| m*Pad4*-R | CTTTGTTCATCTTGGCCTTGG |
| m*Pad2*-F | GAGAAAACCAACTGCGAACTG |
| m*Pad2*-R | CTTCAGGTTTCCATCTCGAGG |
| m*Nox2*-F | TCCTATGTTCCTGTACCTTTGTG |
| m*Nox2*-R | GTCCCACCTCCATCTTGAATC |
| h*PAD4*-F | GGACTGCGAGGATGATGAAG |
| h*PAD4*-R | CACCCTCACTTTGTCCATCTC |
| 18s-F | GTAACCCGTTGAACCCCATT |
| 18s-R | CCATCCAATCGGTAGTAGCG |

**Supplementary Table 3.** Primer details**,** The first lowercase letter of each primer indicates species, including human (h), and mouse (m). Primer names ending with F represent forward primers, while R represent reverse primers.
